# Optical analysis of the action range of glutamate in the neuropil

**DOI:** 10.1101/2021.02.05.429974

**Authors:** E.A. Matthews, W. Sun, S.M. McMahon, M. Doengi, L. Halka, S. Anders, J.A. Müller, P. Steinlein, N. Vana, G. van Dyk, J. Pitsch, A.J. Becker, A. Pfeifer, E.T. Kavalali, A. Lamprecht, C. Henneberger, V. Stein, S. Schoch, D. Dietrich

**Affiliations:** Department of Neurosurgery, University Hospital Bonn, Germany; Section for Translational Epilepsy Research, Department of Neuropathology, University Hospital Bonn, Germany; Institute of Physiology, Medical Faculty, University of Bonn, Germany; Institute of Cellular Neurosciences, Medical Faculty, University of Bonn, Germany; Department of Pharmaceutics, Institute of Pharmacy, University of Bonn; Department of Epileptology, University Hospital Bonn, Germany; Institute of Pharmacology and Toxicology, University Hospital, University of Bonn, Germany; Department of Pharmacology and the Vanderbilt Brain Institute, Vanderbilt University, Nashville, TN, 37240-7933, USA; German Center for Neurodegenerative Diseases (DZNE), Bonn, Germany & Institute of Neurology, University College London, London, United Kingdom

## Abstract

The wiring scheme of neurons is key to the function of the brain. Neurons are structurally wired by synapses and it is a long-held view that most synapses in the CNS are sufficiently isolated to avoid cross-talk to AMPA receptors of neighboring synapses. Here we report in hippocampal brain slices that quantal glutamate release activated optical reporter proteins >1.5 µm distant to the releasing synapse. 2P-glutamate uncaging was used to quantitatively probe glutamate spread in the neuropil. Releasing ∼35000 molecules of glutamate (∼5 vesicles) at a distance of 500 nm to a spine generated an uncaging EPSC reaching ∼30% of the quantal amplitude at synaptic AMPA-Rs. The same stimulus activated ∼70% of the quantal amplitude at NMDA-Rs and still generated clear current and calcium responses when applied at >= 2 µm remote to the spine. Extracellular spread of glutamate on the sub-micrometer scale appeared cooperative and caused supra-additive activation of AMPA-Rs in a spine. These observations are not predicted by previously used models of glutamate diffusion in the neuropil. An extracellular glutamate scavenger system weakly reduced field potential responses but not the quantal amplitude, indicating that a cross-talk component regularly contributes to synaptic transmission. Our data suggest that slight synaptic crosstalk responses at AMPA receptors of ∼2-4 adjacent synapses may be common (>70 synapses for NMDA receptors). Such broadcasting of synaptic signals to very local neighborhoods could stabilize network learning performance and allow for integration of synaptic activity within the extracellular space.

## Introduction

The billions of neurons in the brain are wired to networks and form functional ensembles which are essential to generate distinct behaviors [1]. Neurons are connected structurally and functionally via submicrometer sized junctions, called synapses. At these synapses, neurons signal to the downstream/postsynaptic neurons by releasing diffusible neurotransmitter from presynaptic vesicles. The function of synapses goes beyond simple relay stations and they represent the major element for memory formation and storage of information in the brain by virtue of their adjustable synaptic strength [2-4]. The storage capacity of the brain scales with the number of synapses which operate independently [3,4]. To ensure synaptic independence and avoid diffusible neurotransmitter activating neighboring neurons, synaptic junctions are surrounded by astrocytes which take up and clear released neurotransmitter [5]. Synaptic cross-talk and transmitter spillover has been tested extensively in the past and for the largest part of the CNS it is generally accepted that the firing of neuronal ensembles and computations performed by networks arise from point-to-point signaling only amongst synaptically connected neurons [6]. Consequently, current strategies to decipher simple and higher order brain functions define and reconstruct behaviorally relevant neuronal circuits by only considering hard-wired, synaptically connected neurons [7-10]. Mammalian brains are tightly packed with synapses (∼2/µm^3) at an average nearest neighbor distance of only ∼450 nm and those closely spaced synapses mostly originate from different presynaptic neurons [11-14]. Therefore, the question how relevant transmitter spillover is translates into the problem whether transmitter molecules could partially escape the synaptic cleft and uptake mechanisms and activate neurotransmitter receptors at a distance of ≥0.5 µm [11,15]. If that were the case, clusters of neighboring synapses and thereby non-synaptically connected neurons would regularly be co-activated. Surprisingly, the physical distance at which synaptically released glutamate can activate glutamate receptors on adjacent membranes has not yet been experimentally determined [6] and was only amenable to theoretical analysis (see below).

Synaptic transmission via AMPA-type glutamate receptors (AMPA-Rs) mediates the spread of neuronal activity in most of the circuits of the mammalian brain. For more than two decades theoretical biophysical studies concluded that cross-talk to synaptic neighbors is almost impossible as only high micromolar concentrations of glutamate activate AMPA-Rs and synaptically released glutamate would readily be diluted in the extracellular space [11,15-18]. Most glutamatergic synapses also contain NMDA-Rs which are activated by much lower glutamate concentrations. Accordingly, modeling studies predicted a higher probability for NMDA-Rs to respond to cross-talk from neighboring synapses. In agreement with this view, a number of functional studies of cortical synaptic terminals identified cross-talk mediated by NMDA-Rs [19-23] [24,25]. Similar approaches have largely failed to provide evidence for synaptic cross-talk mediated by AMPA-Rs [20,21]. However, cross-talk via AMPA-Rs will be less pronounced and may be much more difficult to detect experimentally due to the receptor’s lower affinity to glutamate. AMPA-R mediated cross-talk will be smaller when compared to that mediated by NMDA-Rs and AMPA-Rs would also respond only to activity from a much smaller neighborhood. Therefore cross-talk mediated by this type of receptor will require a high spatial density of activated synapses which may be difficult to achieve experimentally.

However, synaptic activation of AMPA receptors outside the cleft seems generally possible and could be experimentally shown under certain conditions. Firstly, in brain regions showing a low density of glutamate transporters or when glutamate uptake is blocked pharmacologically AMPA-R are activated in neighboring synapses and at perisynaptic sites, respectively [26,27]. Secondly, synaptic cross-talk at AMPA-Rs could also be demonstrated at certain specialized synapses where the geometry of the synaptic micro-architecture favors glutamate spill-over or where axons form numerous adjacent synaptic boutons [28-36]. However, in all of these studies providing evidence for synaptic cross-talk at NMDA-Rs or AMPA-Rs the physical distance at which cross-talk occurred remained unknown because neither the glutamate-releasing synapses nor the site of receptor activation could directly be localized. This uncertainty not only makes it difficult to assess how glutamate mediated cross-talk may contribute to the activation of neuronal ensembles but also to exclude the possibility that less pronounced AMPA-R mediated cross-talk might be a phenomenon more widespread than anticipated amongst the pervasive cortical synapses.

Here we directly visualized individual active synapses and the spread of synaptically released quanta of glutamate with novel optical reporter proteins. We further created defined point-like, diffraction-limited, sources of glutamate to quantitively probe the action range of glutamate on the submicron scale with 2P-photon glutamate uncaging. Our results suggest glutamate mediated synaptic cross-talk at AMPA and NMDA receptors may be more widespread than hitherto anticipated. Cross-talk responses of up to several pA may be expected to happen following multi-vesicular release of nearby synapses or coincident activity of multiple synapses in an extracellular microdomains. Our results suggest that synaptic independence could be sacrificed in favor of redundancy and a previously unrecognized form of extracellular synaptic integration and that transmitter diffusion in the neuropil on the sub-micron scale may have to be rethought.

## Results

The recently introduced optical glutamate reporter protein iGluSnFr, when expressed on neuronal membranes, provides a unique way to visualize synaptic glutamate signals [37] and we and others have recently shown that it can also be used to detect quantal glutamate release events [38-42]. Here, we employed iGluSnFr to visualize the action range of glutamate following glutamate liberation by presynaptic exocytosis in brain tissue. We virally expressed iGluSnFr (pAAV1/5-hSyn-iGluSnFR) throughout neurons in the CA3 region of the hippocampus (Fig. 1A) to visualize the spread of glutamate. We chose to examine transmitter release from granule cells as their synapses, the mossy fiber boutons can easily be identified in the hilus by 2P microscopy (Fig. 1A). To unequivocally stimulate only a single mossy fiber bouton in the hilus we patch-clamped granule cells and evoked action potentials by intracellular current injection in the presence of the glutamate receptor antagonists CNQX and APV. By loading granule cells with a tracer dye the stimulated axon and its synaptic boutons embedded in iGluSnFr-expressing neuropil could clearly be identified (Fig. 1A). This allowed us to place line scans across activated boutons with a 2P-scanning microscope. We selected small synaptic boutons for recording and did not include giant mossy fiber boutons which are typically found in the stratum lucidum. The synaptic boutons reported here displayed an average diameter of 0.86 ± 0.14 µm. Triggering single action potentials in granule cells produced either a rapid onset fluorescent response occurring immediately after the action potential and at the position of the dye-filled bouton or a failure as would be expected from the stochastic nature of synaptic vesicle release (Fig. 1B, C). These signals peaked at 27.0 ± 3.4 % and quickly decayed back to baseline (τ = 69 ± 9 ms) but showed a very fast and extended spatial spread. The spatial width of the iGluSnFr signals exceeded the dimension of the bouton (Fig. 1B, D) by several-fold. When averaging the line scans acquired during ‘response’-trials from an individual bouton (Fig. 1D, E) the resulting image displayed an excellent signal-to-noise ratio and the spatial spread of synaptically released glutamate could be directly quantified as a gradual decrease in fluorescent peak amplitude with distance from the stimulated synapse (Fig. 1D-F). Even at a distance of ≥1.5 µm from the bouton a “peak-shaped” fluorescent response could clearly be observed following an action potential (Fig. 1E) which, at that distance, is not expected based on established models of glutamate diffusion in the neuropil e.g. [11,15] (see discussion). The peak amplitude of these synaptically evoked responses exponentially decayed with distance from the bouton and could be well described by λ_sniff_syn_ being 1.2 ± 0.05 µm (Fig. 1F). Note that λ_sniff_syn_ does not provide a direct and proportional read-out of distance-dependent iGluSnfr activation and that λ_sniff_syn_ therefore cannot be interpreted in the usual way of a length constant, i.e. that iGluSnfr activation at λ_sniff_syn_ equals 1/e of iGluSnfr activation in the synapse (due to sublinear response and optical convolution near the synapse, see discussion). Nevertheless, the exponential fits well described the decay of iGluSnFr amplitudes and we therefore use λ to quantitively describe the apparent extend of the optical signal and to compare distance-dependence of glutamate-induced responses across experiments in this study.

**Figure 1.**
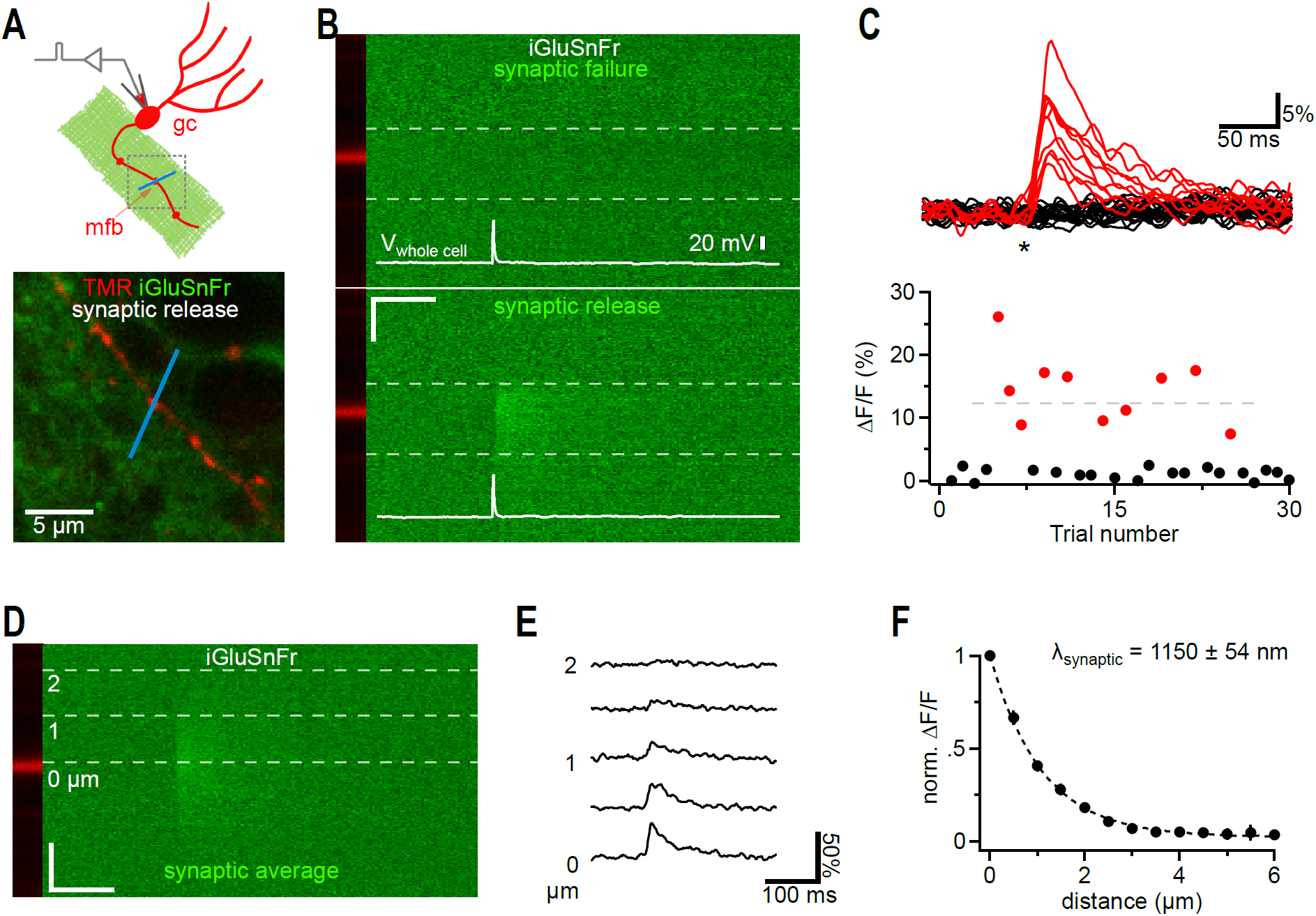
iGluSnFr responses far away from synaptic release sites. (A) Cartoon illustrating the recording condition to quantify the spatial spread of synaptically released glutamate. A granule cell (gc) was patch clamped and dye-filled (red) to identify a synaptic bouton (mossy fiber bouton, mfb) surrounded by neuronal iGluSnFr expression (green). To visualize synaptic glutamate release, a 2P line scan was drawn through the bouton (blue line). The boxed region represents a typical frame scan (as illustrated in lower panel) obtained to identify boutons and adjust the line scan. Lower panel, example dual channel two photon frame scan of a dye-filled (TMR 400 µM, red, iGluSnfr, green) mfb used for stimulation and recording of synaptic glutamate release (as shown in I-L). The bath solution contained CNQX (10 µM) to eliminate network activity and 4-AP (100 µM) and DPCPX (1 µM) to elevate release probability and thereby shorten the required recording time (release probability normally below 10%). (B) Dual channel 2P line scan through the bouton shown in A). Top panel shows a failure, bottom panel successful glutamate release, respectively. White lines illustrate the corresponding whole cell current clamp recordings of the stimulated action potentials. The bouton is located in the red channel displayed on the left and does not show changes in fluorescence (tracer dye). In each line scan image, the region between the two dashed grey lines was used to calculate the fluorescence over time traces shown in C). Note the rapidly rising signal only occurring at the position of the bouton and at the time of the action potential. Scale bar: 50 ms, 1 µm. (C) Top panel, 30 example traces of line scan fluorescence over time (as indicated in B)) demonstrate the well-known typical fluctuation of responses (red) and failures (black) known from synaptic vesicle release. Asterisk, time of action potential. Fluorescence normalized to pre-stimulus levels. Bottom panel, peak amplitudes of the fluorescence traces for the 30 sequential stimulations obtained from this bouton. Markers below the dashed horizontal line represent events putatively classified as single vesicle release responses. (D) Line scans of synaptic responses only (excluding release failures) were averaged per bouton to improve signal-to-noise ratio for quantification of the spread of synaptically released glutamate. Grey dashed lines indicate distances from the center of release. (E) Fluorescence over time extracted from D) at the indicated distances. Note that the decay is slowed with distance and that weak signals can still be detected at 2 µm. The peak of these signals was quantified and plotted in F). (F) Synaptically released glutamate activated iGluSnFr at distances of more than 1.5 µm (n=6). In each experiment the peak amplitudes of fluorescent transients were normalized to the largest amplitude measured at the dye-filled bouton.

Even relatively small synaptic boutons such as those analyzed here may release multiple vesicles in response to single action potentials [39,43-46] consistent with the large variation in iGluSnFr response amplitudes observed (Fig. 1C). A larger amount of glutamate released, by multiple vesicles, may favor spread into the extracellular space [11,15]. To address how much the apparent spread of GluSnFr signals depends on the amount of glutamate released, we selected only the smallest responses obtained by stimulation (e.g. the smallest 4 red dots below the dashed line of the experiment in Fig. 1C) for averaging and analysis (Fig. 2A, B). Despite their lower amplitude (12.7 ± 1.4 %, τ = 71.3 ± 17.5 ms) this subset of responses still showed a comparable λ_sniff_syn_ of 1.4 ± 0.07 µm (n=15, Fig. 2A).

**Figure 2.**
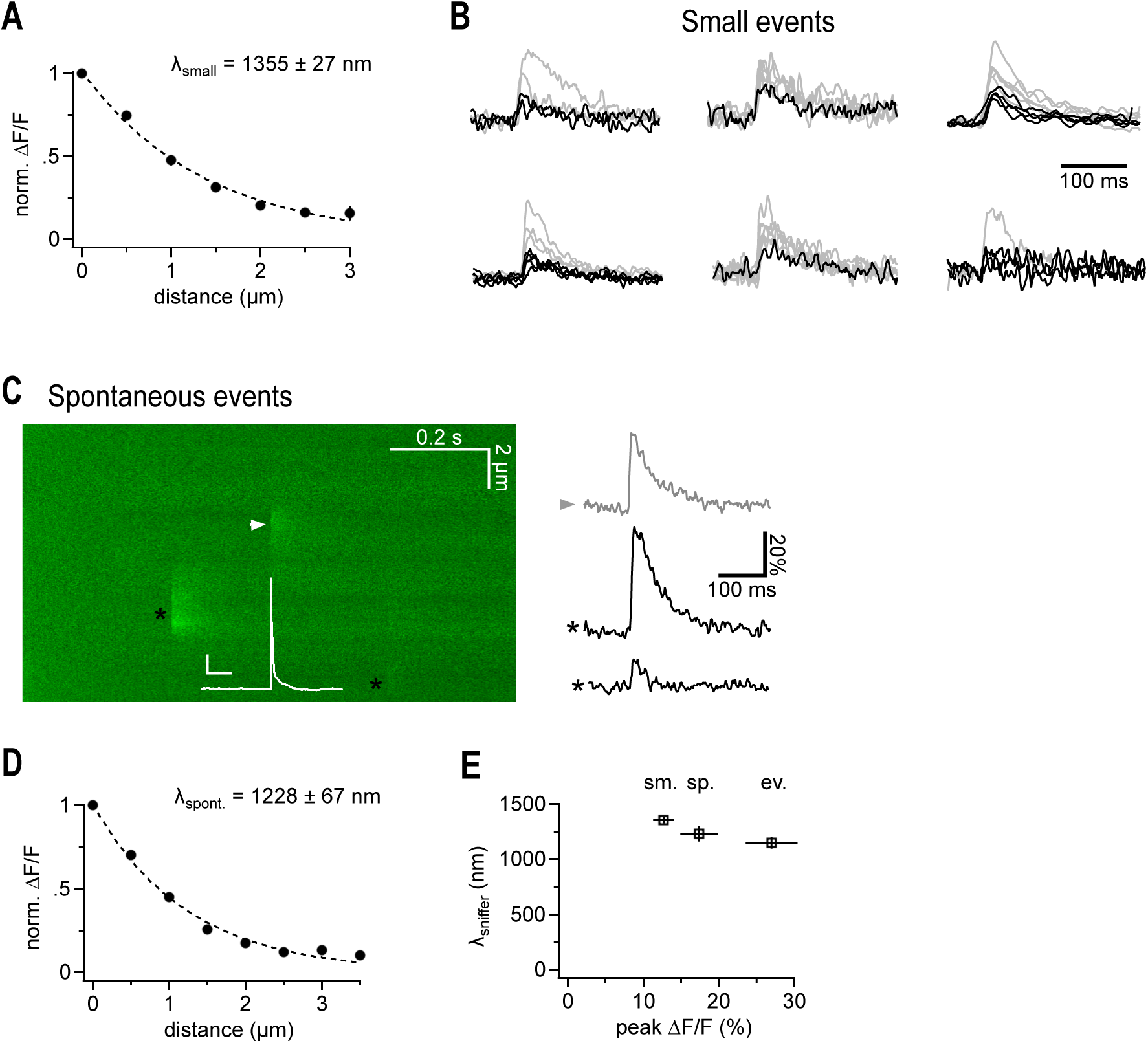
Putative quantal iGluSnFr signal show a similarly extended spatial decay. (A)Spatial extent of small synaptic iGluSnFr transients. For each recording the smallest events were selected to exclude potential multi-quantal events. Note that the lambda value is in the same range as the one derived from data obtained by averaging small and large transients (cf. Fig. 1). (B) iGluSnFr fluorescent traces, the selected fraction of traces used for A) is shown in black. Traces are peak-scaled for comparison are not shown at the same vertical scaling. Events for the cell at the right top are shown in Fig. 1. (C) Example 2P line scan across a dye filled bouton showing the action potential-elicited iGluSnFr response (arrowhead), and 2 spontaneous, off-bouton events (black asterisks) used for the analysis shown in D). The white trace represents the simultaneous current clamp recording of the cell stimulated to fire an action potential which released transmitter at the arrow head position (scale bars: 20 mV, 50 ms). Right panel illustrates fluorescent example traces calculated at the positions indicated by the symbols. (D) Spatial extent of spontaneous likely miniature glutamate transients. Events were analyzed that occurred independently of the timing of the action potential induced in the patch-clamped granule cell and all of them must have been released from neighboring synapses because they did not occur at the dye filled bouton. As spontaneous action potential firing of granule cells in slices is very rare and slices are also bathed in CNQX and APV, these events are likely due to miniature, action potential-independent, single vesicle glutamate release. (E) λ_sniffer_ only weakly depends on the magnitude of the signals and tends to be larger if more glutamate is released. From left to right: selected small, spontaneous and evoked events.

During optical iGluSnFr recordings of stimulated mossy fiber boutons we also observed spontaneous fluorescent transients which occurred in the neighborhood to the stimulated bouton and were not correlated to the action potential (Fig. 2C, n=26). As excitatory transmission was blocked, firing of hippocampal neurons is rare and these events likely represent the optical correlate of spontaneous, action potential-independent, single vesicle release events. These miniature iGluSnFr transients displayed an amplitude (17.4 ± 2.5 %) comparable to that of the subset of small stimulated responses and decayed with a very similar time constant (Fig. 2C, τ = 52.1 ± 6.1 ms). The λ_sniff_syn_ of spontaneous transients confirmed the above quantifications and amounted to 1.2 ± 0.07 µm (Fig. 2D). Taken together, these results strongly suggest that synaptically released glutamate can leave the synaptic cleft and spreads sufficiently far into the extracellular space to activate iGluSnFr molecules expressed on membranes of neighboring cells at >1.5 µm. The extent of this spread only weakly depended on the amount of glutamate released (Fig. 2E).

2-photon (2P) based glutamate uncaging (MNI-caged-glutamate) can be used to generate a small and transient source of glutamate in brain tissue that can be optimized to produce postsynaptic receptor activation similar to synaptic responses [47]. While such an uncaging based point-like source of glutamate is clearly of larger size than a synaptic cleft, it holds the advantage that its position and distance to a synapse can systematically be varied. To compare this approach to synaptic release of glutamate we combined virus-based iGluSnFr expression in CA1 with 2P glutamate uncaging (all following experiments were performed in CA1). Uncaging conditions including MNI-caged-glutamate concentration and laser pulse, were fixed for the whole study and set such that when a laser pulse was applied to a spine head it on average produced an uncaging response (uEPSC) of ∼12 pA (V_h_ −65 mV, APV, TTX see below), which is comparable to miniature EPSCs (see methods for details). Under these conditions a single uncaging pulse in the dendritic neuropil generated spot-like iGluSnFr responses (Fig. 3A, CNQX, APV and TTX were included in the bath) showing larger peak amplitudes (0.81 ± 0.04 DF/F, n=22) when compared to synaptic signals, but rise and decay kinetics were maintained (cf pink trace). Uncaging-induced iGluSnFr fluorescence profiles yielded an only slightly larger λ_sniff_unc_ than the synaptic counterpart (∼1.5 µm, Fig. 3B). The uncaging technique allowed us to test isotropy of glutamate diffusion on the micron scale. For this, lines were scanned through the uncaging spot either perpendicular or in parallel to axons to test for a potential micro-anisotropy of glutamate diffusion in the extracellular space. However, the peak of the fluorescent signals decayed with a very similar length constant when probed parallel or perpendicular to axons (Fig. 3B). Lack of anisotropy of glutamate diffusion was confirmed by using long (250 ms) iontophoretic applications of glutamate which produced similar near steady state spatial gradients of glutamate in both orientations (Suppl. Fig. 1).

**Figure 3.**
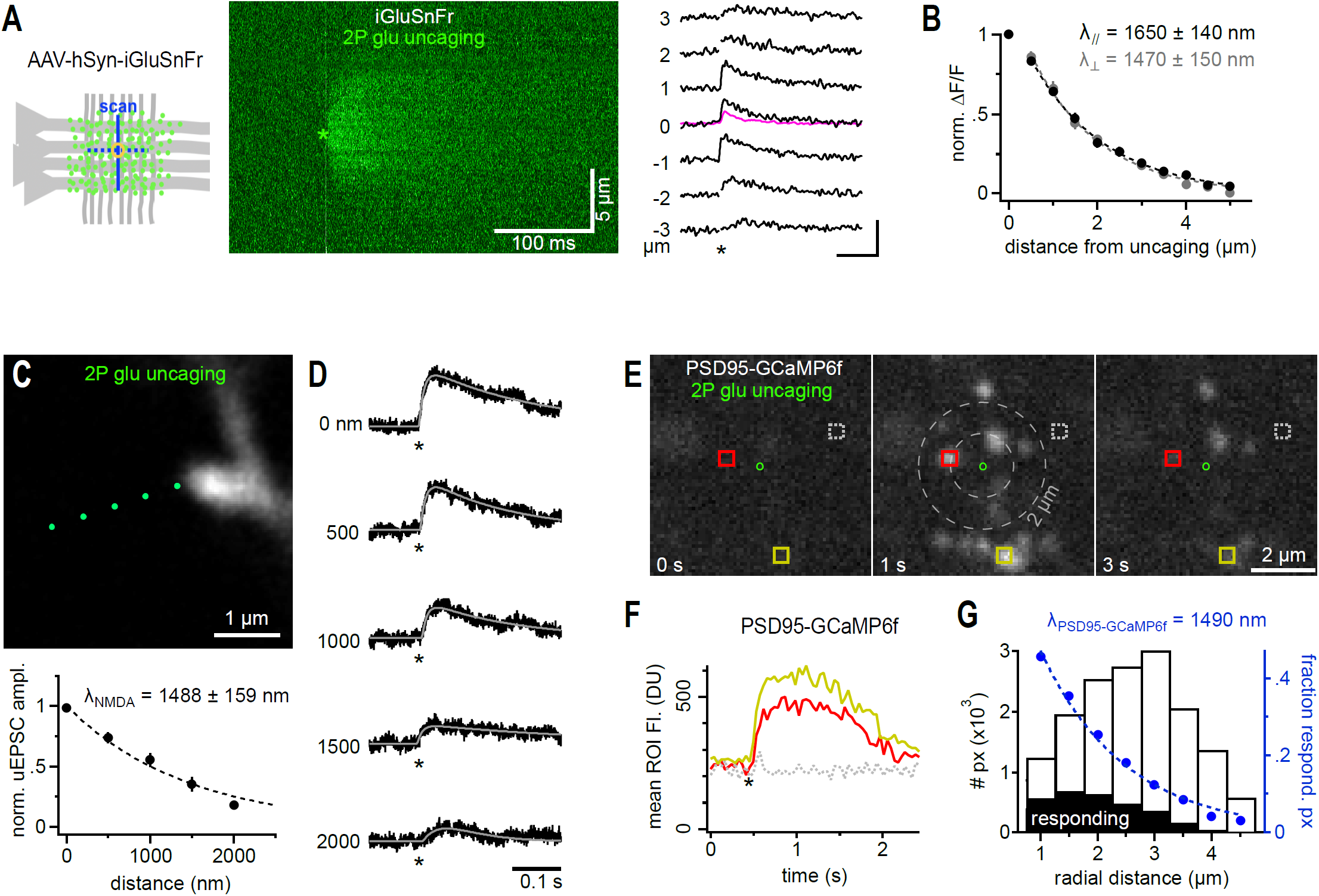
Extended spatial decay of NMDA-Rs mediated signals. (A)iGluSnFr reports a similar spread of extracellular glutamate following 2P-glutamate uncaging. Cartoon: yellow circle indicates glutamate uncaging site in the dendritic region of CA1 (Str. radiatum) where iGluSnFr reporter proteins are expressed on neuronal membrane (green dots). 2P line scans perpendicular or parallel to axons (blue lines) were used to quantify the spatial spread of the fluorescent signal. Middle panel: Example line scans through the glutamate uncaging site (green asterisk, indicating time and position, average of 3 repeated uncaging spots at 3 s intervals). Note the rapid and substantial spread of the fluorescence. Line scans were normalized on the pre-uncaging fluorescence to account for spatial variability of initial iGluSnFr brightness (owing to varying spatial densities of membrane expression levels). Right panel: example fluorescent traces calculated from the line scan image shown in the middle. Numbers indicate distance from uncaging site; asterisk, time of uncaging. Note the visible and delayed signal at ± 3 µm. Kinetics and amplitude are similar to synaptically evoked iGluSnFr responses as illustrated by the pink trace, average response from the experiment shown in Fig. 1. Scale bar: 100 ms, 100% (B) λ_sniff_unc_ measured from iGluSnfr signals is isotropic (n=10 for each direction, black and grey markers represent scans parallel and perpendicular to axons, respectively) and only slightly exceeds λ_sniff_syn_ obtained following synaptic glutamate release. (C) 2P scan of a dye-filled spine incubated in 20 µM CNQX and 1 µM TTX to isolate NMDA receptors. Uncaging spots (green) were separated by 500 nm and applied at 5 s intervals to account for the substantially slower kinetics of NMDA-R mediated uEPSC. Lower panel: λ_NMDA_ after glutamate uncaging (n=12). (D) Example traces of NMDA receptor-mediated uEPSCs (asterisk, time of uncaging, cell voltage clamped at +40 mV). uEPSCs are still clearly seen at a distance of 2 µm and their kinetics are substantially slower. To reliably quantify peak amplitudes of even the smallest responses uEPSCs were fitted with a two-exponential function (grey line, see methods). Note that even remotely evoked uEPSCs (>1500 nm) evoke clear currents demonstrating pronounced diffusional propagation of released glutamate. (E) Widespread activation of PSD95-GCamp6F following a single uncaging pulse confirms large action range of glutamate at NMDA receptors. Three 2-photon scans (take from the 20 Hz time series quantified in F)) in the dendritic region of CA1 before and after the uncaging pulse (green circle indicates uncaging site, 15 µM glycine to allow NMDA-R activation at resting potential, 20 µM CNQX, 1 µM TTX). Note the appearance of bright spine head-shaped structures following glutamate uncaging which occur even outside a 2 µm range (grey dashed circles). Colored squares indicate example ROIs used to calculate the fluorescence over time traces displayed in F). (F) Average ROI fluorescence over time illustrating the pronounced calcium increases induced in spine heads by activation of NMDA receptors following glutamate uncaging (asterisk, colors of traces correspond to the ROIs shown in E). (G)Estimation of λ_NMDA_ from the spatial distribution of calcium responses (PSD95-GCamp6F) around the uncaging point. The histogram plots the frequency of responding pixels (see methods for threshold details) along the radial distance from the uncaging site (black bars, “responding”, aggregated results over 66 uncaging events). The white bars show the number of pixels in the acquired image along the radial distance. The ratio of the black over the white bars represents the experimental probability of observing a calcium response at a given distance (blue markers, fraction of responding pixels). This probability drops with distance and follows a λ_NMDA_GCaMP_. Notably, λ_NMDA_GCaMP_.

Such remote action of glutamate in the extracellular space might also cause physiologically relevant activation of remote glutamate receptors. As iGluSnFr binds glutamate with a similar high affinity as NMDA receptors [37,48] our results predict NMDA receptor activation following uncaging at distances >1.5 µm. As spines are known sites of postsynaptic glutamate receptor clusters [49] we identified spines by patch clamping and dye-filling CA1 pyramidal neurons in hippocampal slices (Fig. 3C). We applied the above described glutamate uncaging protocol and recorded the distance-dependent decay of NMDA receptor-mediated uEPSCs (Fig. 3D). To isolate NMDA receptor currents (uEPSC_NMDA_), we voltage-clamped cells at +40 mV (Cs-based intracellular solution) and blocked AMPA receptors. We targeted those spines for uncaging which lacked neighboring spines of the recorded neuron within a sphere of at least ∼2 µm diameter to minimize the possibility that other spines contribute to the electrical response by binding diffusing glutamate. When we moved the uncaging laser spot away from the spine head, uEPSCs_NMDA_ clearly but slowly declined and were still detectable at a substantial distance of at least 2 µm (Fig. 3C, D). The decay could also be well described by an exponential function with a length constant of 1488 ± 159 nm (n=12. Fig. 3C) and we refer to this length constant as λ_NMDA_. This large range together with the high density of synapses (∼2 µm^-3^) implies that photo-released glutamate should activate NMDA receptors on a multitude of neighboring spines around the uncaging spot. To directly visualize this prediction and show that this activation translates into a physiologically relevant down-stream signal, we virally expressed the genetically encoded calcium indicator GCaMP6f in CA1 pyramidal cells. GCaMP6f was fused to PSD95 which selectively targeted it to dendritic spines (Fig. 3E). Before stimulation, fluorescence of the calcium indicator was very dim and spines were almost invisible. However, a single uncaging pulse of glutamate in the center of the scan field (green circle) produced simultaneous pronounced increases in fluorescence in many spines of transduced neurons around the uncaging spot (Fig. 3E, F). Importantly, not only spines very close to the uncaging site but also those at a distance of >2 µm were activated, consistent with the λ_NMDA_ estimated by uEPSCs_NMDA_ recorded at increasing distances from an individual spine. In fact, when accounting for the geometric, random occurrence of PSD95-GCaMP6f expressing spines the probability of finding a responding spine drops with the distance from the uncaging spot following a length constant, λ_NMDA_GCaMP_, of ∼1.5 µm (Fig. 3G, note that not all neurons were transduced by the virus). Thus, a single uncaging spot of glutamate simultaneously activates NMDA receptors on a very large number of spines from probably at least many tens of neighboring neurons in the 3D environment and receptor activation induced pronounced postsynaptic Ca^2+^ signals.

Most excitatory activity of the brain is transmitted from neuron to neuron by AMPA-Rs. AMPA-Rs show a substantially lower affinity for glutamate than NMDA-R and iGluSnFr. For this reason, AMPA-Rs should produce smaller responses to remote glutamate sources. Here, we used 2P glutamate uncaging to test at which distance synaptic AMPA receptors still respond to glutamate. As mentioned above, delivery of glutamate by brief laser pulses at the spine head produced an uEPSC of 12.4 ± 1.0 pA (Fig. 4A, in the presence of TTX, APV and gabazine, see methods for details on uncaging conditions) comparable to the amplitude of mEPSCs measured in the same cells (11.5 ± 0.6 pA, n = 8). uEPSCs were mediated by AMPA receptors as they were completely blocked by CNQX (Suppl. Fig. 2). When we moved the uncaging laser spot away from the spine head, responses declined much faster than for NMDA receptors but were still clearly detectable at a distance of >600 nm (Fig. 4B). We found λ_AMPA_ to be 450 ± 34 nm (n = 27, Fig. 4C). λ_AMPA_ was similar when probed at different angles to the Schaffer collaterals (not shown) or at shaft synapses (Suppl. Fig. 2) implying that average diffusion and uptake on the sub-micrometer scale are isotropic and a function of the random shape of the extracellular space immediately surrounding synapses (at ∼0.5 µm). This is consistent with the above described results obtained with iGluSnFr, a macroscopic diffusion analysis in this brain region [50], and the finding that the structure of the neuropil surrounding synapses appears random and chaotic [11].

**Figure 4.**
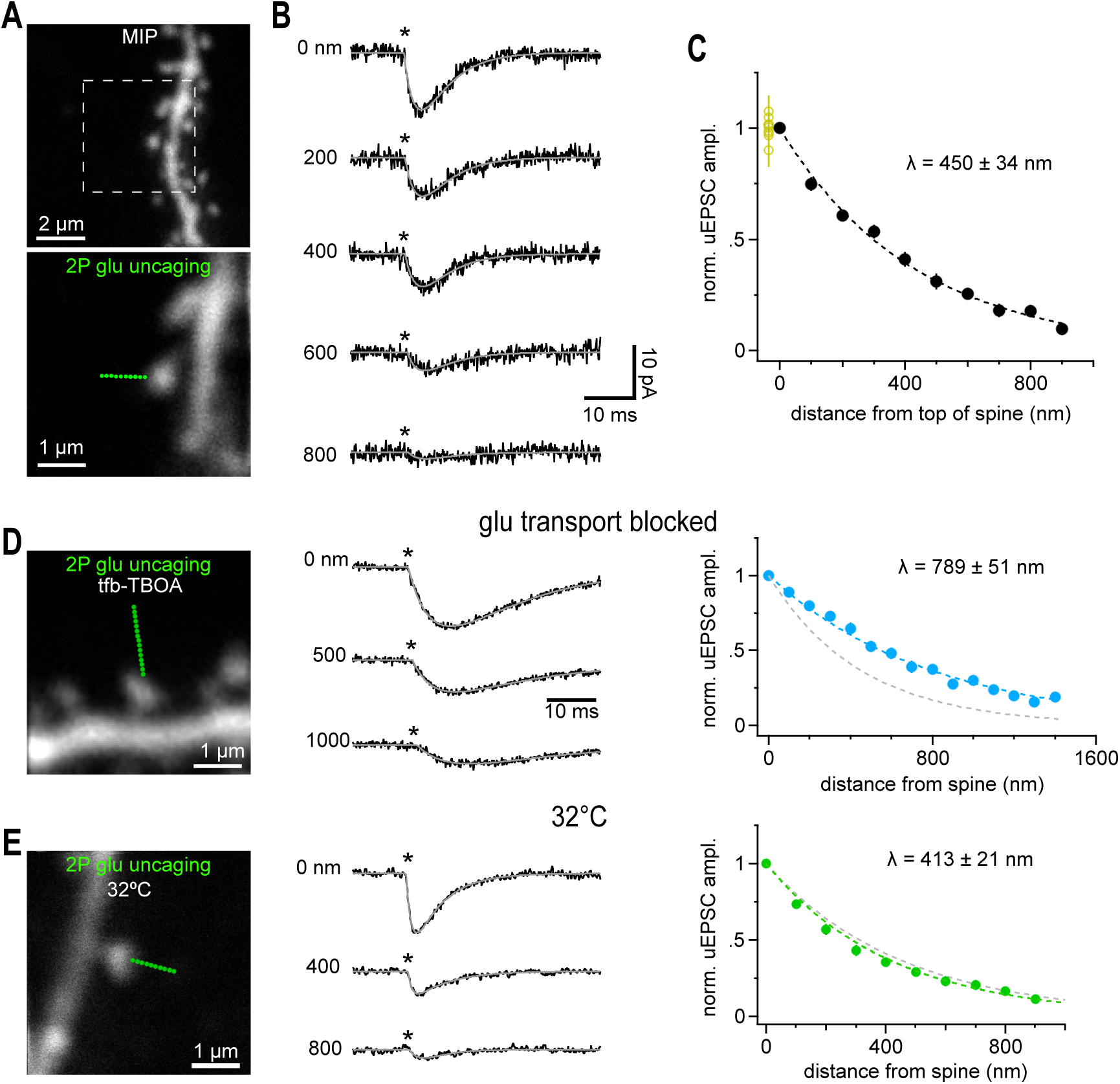
Glutamate uncaging beyond the nearest synaptic neighbor distance activated also activates synaptic AMPA receptors. (A)Maximum intensity projection (MIP) of CA1 pyramidal cell dendrite dialyzed with 25 µM AlexaFluor 594 scanned with a two-photon microscope. Solitary spines were selected to avoid co-activation of neighboring structures. Lower image illustrates positioning of a sequence of glutamate uncaging points (green dots, step size 100 nm) to probe the spatial dependence of uEPSC amplitudes. Single image scanned at higher resolution. (B) Example current traces recorded in whole-cell voltage clamp mode showing the gradual decline of the response magnitude with distance. Light pulses (0.6 ms, asterisks) were applied at 1 Hz. Grey lines show fitted with a two-exponential function used to determine the peak amplitude. Note that even uEPSCs evoked at >400 nm peak within ∼3-4 ms reflecting the rapid diffusional propagation of glutamate. Throughout the study we used the following conditions for isolating AMPA-Rs: 720 nm, 0.6 ms, 23 mW, 5 mM MNI-caged glutamate in presence of 1 µM TTX, 50 µM APV, 10 µM Gabazine. (C) Summary graph of the distance-dependent decay of the amplitude of uEPSCs (n=27 spines) which could be well approximated by an exponential function with a length constant λ (dashed black line). Fitting of the individual amplitudes over distance revealed the indicated average value of λ. Applying 10 identical glutamate uncaging pulses at 1 Hz at the spine head yielded stable responses (yellow circles at 0 nm, n=8) indicating that desensitization or run-down of receptors is negligible. (D) Left: 2P-photon scan of a spine incubated in 1 µM tfb-TBOA, 100 µM APV, 40 µM MK801, 10 µM gabazine and 1 µM TTX. Uncaging responses were probed over an extended distance by additional uncaging spots (green dots, step size 100 nm). Middle: Example uEPSCs (averages) taken from the three distances indicated. Note the prominent residual current at 1000 nm (compare to B). Asterisk, time of uncaging pulse; grey line, uEPSC fit. Right: Extended action range of uncaged glutamate in the presence of tfb-TBOA. Blue markers represent the average decay of uEPSCs measured from 21 spines yielding an average λ as indicated. Dashed grey line shows the control λ (450 nm) as determined in C). (E) as in A-C) but slices were kept at 32°C. Compared to results obtained at room temperature the action range of uncaged glutamate at AMPA-Rs is only slightly shortened at 32°C suggesting that transmembraneous transport of glutamate (highly temperature dependent) is too slow to modify extracellular glutamate signalling on this short spatial scale. 32 spines yielded the average λ as indicated. Grey dashed line shows the control λ (450 nm) at room temperature (cf C).

Previous work demonstrated a clear role of largely astroglial glutamate uptake in limiting the spread of glutamate in the extracellular space but how uptake affects the distance at which glutamate can still activate receptors remained not exactly known [18-20,22-24,51]. We found that the λ_AMPA_ was clearly increased ∼1.8-fold by strongly blocking transporters with tfb-TBOA (Fig. 4D, 789 ± 51 nm, n=21, a milder block of transporters by DL-TBOA (∼3000-fold lower affinity at EAAT1) increased λ_AMPA_ to a weaker extent, Suppl. Fig. 2). The glutamate turnover rate (# of glutamate molecules translocated intracellularly per time) is known to strongly increase with temperature [52]. However, λ_AMPA_ probed by uncaging at near-body temperature (32° C) only modestly decreased by ∼15% to 413 ± 21 nm, suggesting that glutamate turnover is not critically involved in the effect of transporters on this submicrometer scale (n=32, Fig. 4E, compared to 450 nm at RT).

During development [53] and in disease [54] glutamate transporter activity or expression levels substantially change. These observations prompted us to test for alterations of λ_AMPA_. While the unchanged time course of synaptically evoked glutamate transporter currents in astrocytes of older mice has been taken as evidence that net extracellular glutamate handling is preserved during development when analyzing bulk synaptic signals [53], we found λ_AMPA_ to be significantly changed on the sub-micron scale around individual spines. λ_AMPA_ in adult hippocampal tissue (5-7 weeks) was ∼25% shorter when compared to the juvenile value (Fig. 5A, 345 ± 20 n, n=39 vs 450 ± 34 nm at P17).

**Figure 5.**
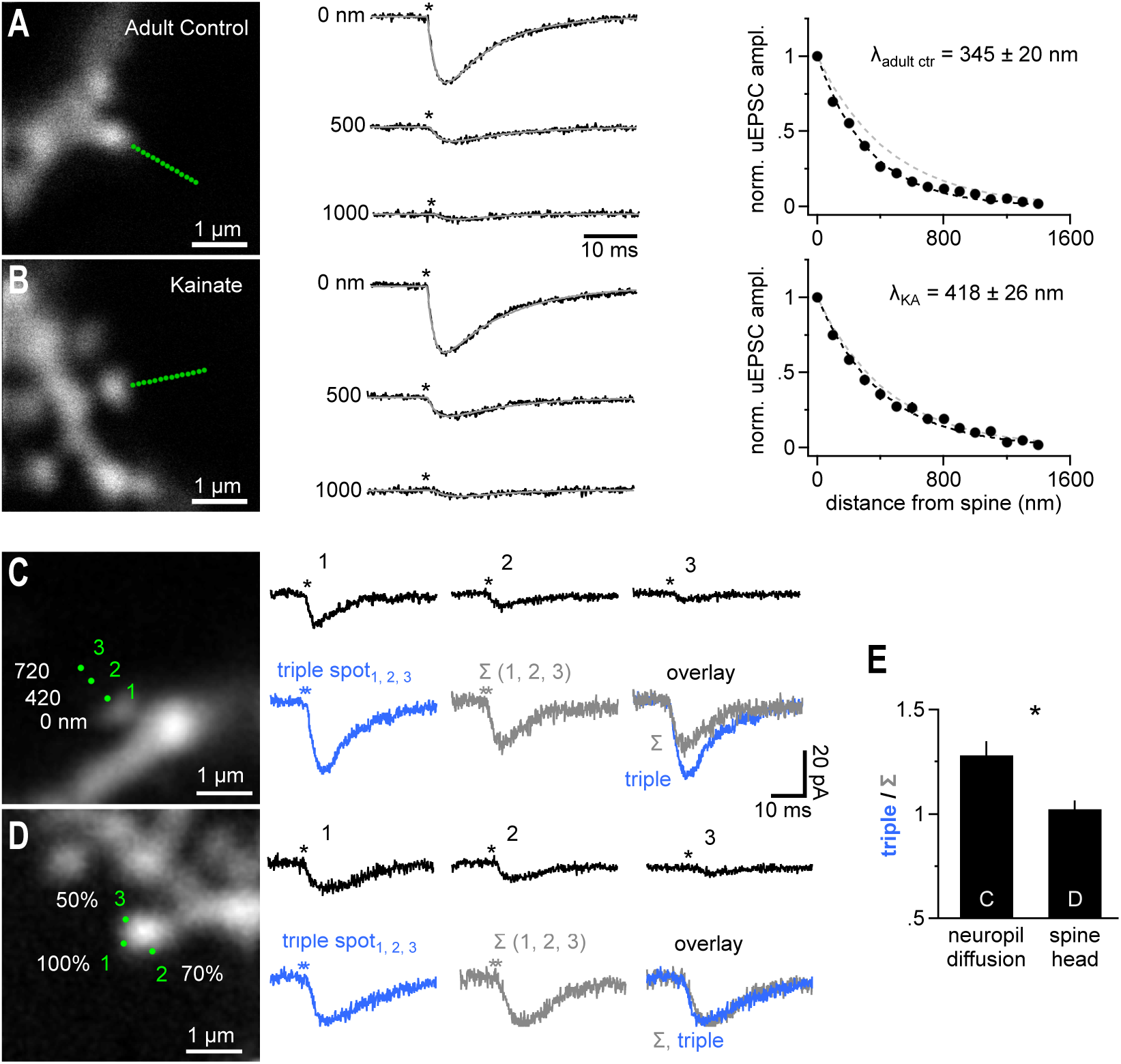
Extracellular temporal integration of glutamatergic released in sub-micron peri-synaptic neighborhood. (A)Adult mice (6 weeks) show a reduced glutamate action range at AMPA receptors (n=39). Conditions as in Fig. 4A-C. Traces represent averages of n=39 recordings. For comparison the right panel also shows the λ determined for adolescent mice (grey dashed line). (B) In an animal model of chronic temporal lobe epilepsy (suprahippocampal kainic acid, injection-induced status epilepticus, n=15) there is a significant extension of the action range of uncaged glutamate at AMPA receptors back to levels seen in adolescent mice (n=25, p=0.024, studentized bootstrap test for difference in means). Note that the two fits (black and grey (450 nm control group, Fig.4A-C) dashed lines) are almost indistinguishable. (C) Left panel, dye-filled spine used to probe the summation of coincident activity in the spine-surrounding extracellular space. 3 uncaging spots (1, 2, 3) were applied at the 3 distances indicated. Right panel, top row shows the responses when the 3 uncaging spots were applied sequentially (asterisks, time of uncaging pulse). Bottom row shows the response to synchronous uncaging at the 3 spots (left, blue, “triple spot”) in comparison to an arithmetic sum (middle, Σ (1, 2, 3)) of the 3 responses shown in the top row. The overlay on the right shows that the triple response clearly exceeds the arithmetic sum. (D) As in C) but the 3 spots were all placed right at the spine head. To mimic the weaker response obtained by uncaging spot 2 and 3 in C) the uncaging laser power per spot was reduced to 70 and 50 %. Scaling as in C). Right panel, top row shows the responses when the 3 uncaging spots were applied sequentially. Bottom row shows the response to synchronous uncaging at the 3 spots (“triple spot”, 100%, 70% and 50% of laser power as in top row) in comparison to an arithmetic sum (Σ (1, 2, 3)). The overlay on the right clearly shows equal response amplitude indicating that the supra-additive summation is not a function of the spine or dendrite. (E) Summary of n = 11 (panel C) and n = 34 (panel D) experiments demonstrating a significantly larger response when uncaging spots were distributed in the neuropil and involved diffusion in the extracellular space.

Glutamate transporters were shown to be down regulated in the early phase of a mouse epilepsy model [54]. Using the same epilepsy model (suprahippocampal kainic acid injections to induce status epilepticus, see methods) in adult mice (6 weeks) we found λ_AMPA_ tested on spines of CA1 pyramidal cells prepared 5 days post injection from contralateral hemispheres to be significantly increased by ∼20% (418 ± 26 nm, n = 25) compared to the adult control group (Fig. 5B). Thus, λ_AMPA_ is not a biological constant but is sensitive to developmental and pathological changes in the extracellular micro-environment of a spine.

If the spread of glutamate in the extracellular space is limited by glutamate transporters, as suggested by the effect of TBOA (cf Fig. 4, see discussion), then coincident glutamate release from nearby sources may cooperate to consume free transporter binding sites and show an enhanced spatial spread. We therefore tested the capabilities of AMPA receptors on spine heads to integrate input from remote sources in the presence of the NMDA receptor antagonist APV (and TTX). We first recorded uEPSCs as responses to three independent, consecutive (1 s interval) uncaging stimuli at three distances from the spine head (Fig. 5C, at 0, 420, 720 nm) and observed a decline with distance consistent with λ_AMPA_ determined above (Fig. 5C, top row). We then re-applied the 3 uncaging pulses at the same positions but this time almost simultaneously (Fig. 5, “triple spot”, see methods). The resulting compound response was clearly larger than each individual uncaging response as expected because we overall released more glutamate. Unexpectedly, the amplitude of this compound uEPSC (“triple spot”) also significantly exceeded the amplitude of the arithmetic sum of the three consecutively acquired uEPSC (“Σ (1, 2, 3)”). Here, the amount of glutamate uncaged is identical and the increase in amplitude suggested an enhanced spread of coincident glutamate release activity. An alternative explanation for this supra-additivity is that the larger electrical signal in response to the triple uncaging stimulation triggers stronger electrical signaling within the spine which may boost the recorded current response (e.g. by recruiting voltage-gated channels). To address this possibility, we re-designed the experiment and positioned all three spots at the spine head to directly probe the responsiveness of spines circumventing diffusion of glutamate (Fig. 5D). The laser power for the second and third uncaging spot was reduced so that the resulting uEPSCs mimicked the size of the responses to uncaging at 420 and 720 nm, respectively and we achieved an equivalent electrical signal. Thus, in this re-designed experiment the degree of glutamate receptor opening in the spine was maintained compared to the original experiment but no diffusion to the spine head is involved. As above, the three laser pulses were first applied consecutively (upper row) and then simultaneously (“triple spot”). In this case, supra-additivity was absent and the amplitude of the simultaneously applied uncaging spots almost exactly equaled the amplitude of the arithmetic addition of the three single uEPSC traces (“Σ (1, 2, 3)”, Fig. 5D, E). This showed that triggering of postsynaptic electrical signaling does not explain the supra-additive summation. Therefore, the supra-additive summation happened in the extracellular space and likely involves facilitated spread of glutamate from the remote spots to the spine head.

This result opens the question of how dependent λ_AMPA_ is on the amount of glutamate being released. Uncaging might release many more glutamate molecules than synaptic release, prompting us to check whether shorter estimates of λ_AMPA_ result if we reduce the amount of glutamate released. We tested the dependence of λ_AMPA_ on the amount of glutamate being released by systematically measuring λ_AMPA_ at individual spines with low, normal and high uncaging laser power changing the free glutamate concentration to ∼64%, 100% and ∼144%, respectively (by altering laser power to 80% and 120%). The amplitudes of the resulting uEPSCs clearly varied with the amount of glutamate released (Fig. 6A). In contrast, λ_AMPA_ did not become shorter when we released less glutamate indicating that λ_AMPA_ is not steeply dependent on the amount of glutamate released (Fig. 6B, C). On the other hand, the λ_AMPA_ was slightly enlarged when we released more glutamate suggesting that our standard conditions generate a glutamate load at the upper end of the extracellular glutamate handling capacity (Fig. 6B, C). This is consistent with the view that when locally exceeding a critical extracellular glutamate concentration, consumption of glutamate binding sites can facilitate the spread of glutamate.

**Figure 6.**
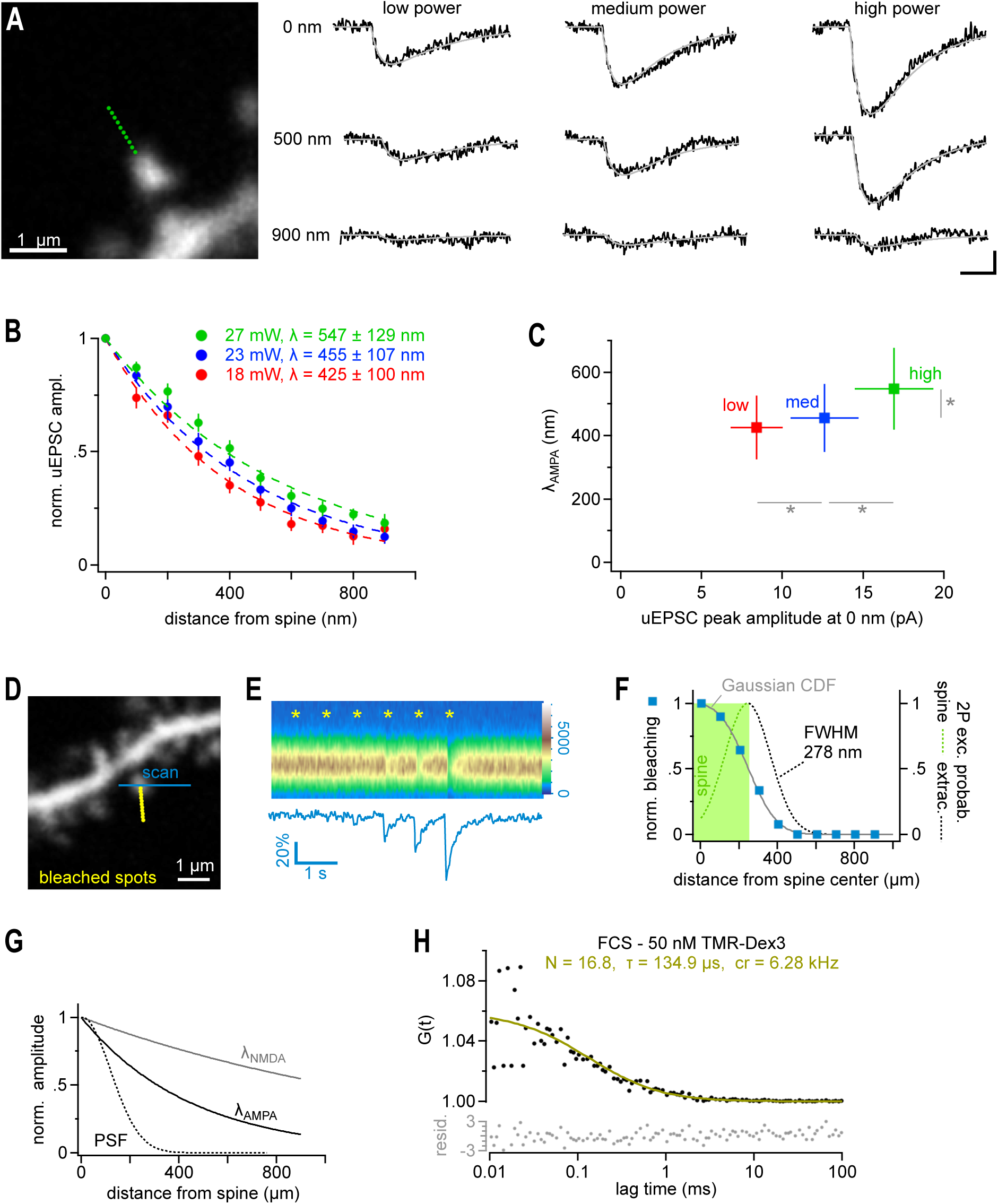
2P-glutamate uncaging does not overwhelm transporters and mimics multi-vesicular release. (A)In order to test the dependence of λ on the amount of glutamate being released by uncaging 10 points were placed from 0 to 900 nm from the edge of the spine head (left panel, green dots). Uncaging power at the objective was set at 18, 23 or 27 mW (changing the free glutamate concentration to ∼61%, 100% and ∼137%, respectively, due to the two-photon immanent non-linear, quadratic, dependence of the uncaging rate on the laser power). All three power levels were tested at each of the 19 spines in a randomized order. Single responses from a representative spine are shown in black, with their double exponential fits in grey (right panel). (B) λ at AMPA-Rs at low (red), medium (blue) and high laser power (green) extracted from all spines recorded as in (A). (C) λ values plotted against the peak amplitude of the uncaging currents recording when uncaging at 0 nm. Note that while the amplitude of the uEPSCs significantly varied with laser power (horizontal grey bars and asterisks, Repeated measures ANOVA, Tukey post-hoc), λ did not decrease when releasing fewer molecules of glutamate despite a significant reduction in uEPSC amplitude suggesting that transporters are not overwhelmed and AMPA-Rs are operating in a near linear range (Repeated measures ANOVA, Tukey post-hoc). In contrast, releasing more glutamate did lead to a significantly increased in the length constant λ (vertical grey bar and asterisk, Repeated measures ANOVA, Tukey HSD post-hoc) indicating that at higher glutamate concentrations further signs of transporter saturation can be observed. Also note that amplitude varies linearly with the estimated amount of uncaged glutamate, also suggesting that AMPA-Rs are operating in a near linear range. (D) Dye-filled dendrite with spines used to probe the optical resolution of our uncaging system in brain slices. The imaging scanner was used to monitor the fluorescent emission from a single spine (blue line, 820 nm). The uncaging laser (720 nm) produced a series of light spots as for uncaging (Δx 100 nm) but the closest spot was placed directly onto the spine head (yellow dots) to measure the maximal bleaching amplitude of the spine with the line scans of the imaging laser. (E) Bleaching amplitude steeply drops off with distance from the spine. Top panel: Repetitive line scans through the spine head (color legend on the right edge). Asterisks indicate the times when bleaching spots were applied. Bleaching spots were sequentially moved towards the spine. Note that bleaching is clearly seen only with the third from last spot (200 nm) due to the small size of the bleaching spot generated by the uncaging laser. Bottom panel: average fluorescence of the scanned lines used to quantify the bleaching amplitudes. (F) Summary graph of 6 experiments as illustrated in D and E to extract an estimate of the full width at half maximum (FWHM) of the diffraction limited spot of the uncaging laser. Normalized bleaching amplitudes extracted from line scans (as in E) are shown as blue squares. As the dimension of our detector of bleaching, the spine volume, is much larger than the bleaching spots (as opposed to the typically used sub-resolution-sized beads typically used in in vitro measurements) the blue squares do not directly yield the spatial resolution or point spread function (PSF). To illustrate this relation the green area shows the volume occupied by a spine and the obtained bleaching is half maximal then, when the PSF is centered on the edge of the spine (dashed line). Then, only the left half of the PSF bleaches the spine (green dash) whereas its right half hits the extracellular space (grey dash). Only once the PSF is fully contained in the spine volume, maximal bleaching is achieved. Therefore, the distance-dependent bleaching amplitudes (blue squares) provide the integral of the PSF (green area under the PSF curve) and have to be fitted by a Gaussian cumulative distribution function (CDF, “integral of Gaussian”, grey line) to extract the approximated shape of uncaging system’s PSF (dashed line) and the FWHM. This analysis suggests, the optical resolution of our uncaging system was close to the theoretical optimum, FWHM = 278 nm (dashed line). (G)Comparison of the estimated PSF (as in F) to the λ values at AMPA-Rs and NMDA-Rs, respectively. Note that the latter two clear exceed the optical resolution of our uncaging system. (H)Fluorescence correlation spectroscopy approach to estimate the 2P-uncaging volume. Fluctuations in emission of a 50 nM TMR-dextran3kD solution during exposure to the stationary uncaging laser beam (720 nm, NA 1) was recorded for 120 s and used to calculate the autocorrelogram (black dots). Fitting the autocorrelogram with an autocorrelation function assuming a 3D gaussian volume (yellow) yielded 16.8 diffusing dye molecules in the effective detection volume. Together with the known dye concentration this estimates the excitation volume to be ∼0.2 fl (including γ-factor correction, see methods for details). Lower panel shows the residuals of the fit (see methods for details of residuals).

To define this critical level of glutamate and relate it to the density of synapses and their activity, the amount of glutamate released during uncaging and its spatial distribution needs to be known. For this reason, we determined estimates of the dimensions of the uncaging 2P point spread function and the number of uncaged glutamate molecules. We assessed the point-spread function of our 2P-uncaging system in situ by monitoring the degree of bleaching of dye-filled spines when we moved the uncaging laser progressively closer to the spine. We selected spines in the slice depth of ∼30 µm, which we also used for uncaging experiments. This allowed us to estimate that the point-spread function of our uncaging laser beam shows a FWHM of ∼278 nm (Fig. 6D-F and see methods). Thus, receptors on a spine head in the focus of our 2P uncaging laser are initially exposed to a gaussian shaped spatial profile of glutamate concentrations with a FWHM of ∼280 nm and the optical resolution of uncaging compared well against λ_AMPA_, λ_NMDA_, λ_NMDA_GCaMP_ and λ_sniff_unc_ (Fig. 6G).

The amount of glutamate molecules released by uncaging increases with the focal excitation volume of our system, the volume in which caged glutamate is converted. We experimentally determined the excitation volume of our system with fluorescent correlation spectroscopy (FCS, see methods for details) to be ∼0.2 fl (Fig. 6H), which is in good agreement with theoretical predictions [55]. The neurons in the slice are bathed in 5 mM MNI-glutamate during the experiment. The number *n* of glutamate molecules released by our uncaging pulse can then be estimated by: *n* = 0.2 fl * 5 mM * ε * ExVF * N_Av_ ∼ 36000, with ExVF being the extracellular volume fraction (0.2) and ε the estimated fraction of glutamate uncaged (0.3, see discussion). Assuming recent estimates for the number of glutamate molecules per synaptic vesicle, ∼7000-8000 [56,57], this calculation shows that our uncaging releases approximately the same number of glutamate molecules contained in 5 synaptic vesicles. We combined those estimates and calculated the approximate glutamate concentration profile around the uncaging position and superimposed it on an electron micrograph of a cortical synapse for a rough spatial comparison (Fig. 6I).

The mean distance to the nearest neighbor synapse in the CA1 region has been reported to be ∼450nm [11]. Photo-releasing ∼36000 molecules of glutamate or the equivalent to ∼5 synaptic vesicles at this distance produced ∼38% of the uni-quantal AMPA-R-mediated amplitude (∼4.6 pA versus 12 pA quantal amplitude). Multi-vesicular release of 2-5 vesicles is a common scenario at hippocampal synapses [39,43-46]. Thus, if our glutamate uncaging responses at this distance mimic multi-vesicular release (see discussion), there should be a small but consistent degree of cross-talk between neighboring synapses. To discern and eliminate such potential cross-talk component we applied a high concentration of glutamate-pyruvate transaminase (GPT) and pyruvate as a biochemical glutamate scavenger system (cf [58], see methods) to inactivate synaptically released glutamate before it reaches a neighboring synapse. If cross-talk is present then GPT application should reduce fEPSPs and this reduction should be even stronger for the second, paired-pulse (40 ms) fEPSP as this recruits a higher spatial density of active synapses due to presynaptic facilitation. Indeed, as shown in Fig. 7A, GPT slightly reduced the first and the second fEPSPs to 92±5 % and 88±4% (n=9), respectively. A similar reduction of synaptic transmission by this scavenger system was observed by [58]. However, the authors attributed it to an effect of GPT on glutamate still in the synaptic cleft, during diffusion to postsynaptic receptors. To test this assumption that GPT can capture glutamate while still in the synaptic cleft we recorded mEPSCs in dissociated neuronal cultures and quantified the mEPSC amplitude, the response to release of a single vesicle (Fig. 7B). If GPT acts in the synaptic cleft it should reduce the mEPSC amplitude. However, GPT did not reduce the amplitudes of mEPSCs indicating that the scavenger system is not potent enough to interfere with intra-cleft glutamate receptor activation (Fig. 7B). Thus, it appears likely that the scavenger indeed reduced fEPSPs by interfering with synaptic cross-talk. Cultures were chosen here as they grow at lower densities than neurons in brain tissue and the nearest neighbor distance of synapses typically exceeds 1 µm – a distance at which cross-talk of quantal responses should be minimal and undetectable [59,60]. Therefore, mEPSCs in culture allowed us to record direct, synaptic activation only.

**Figure 7.**
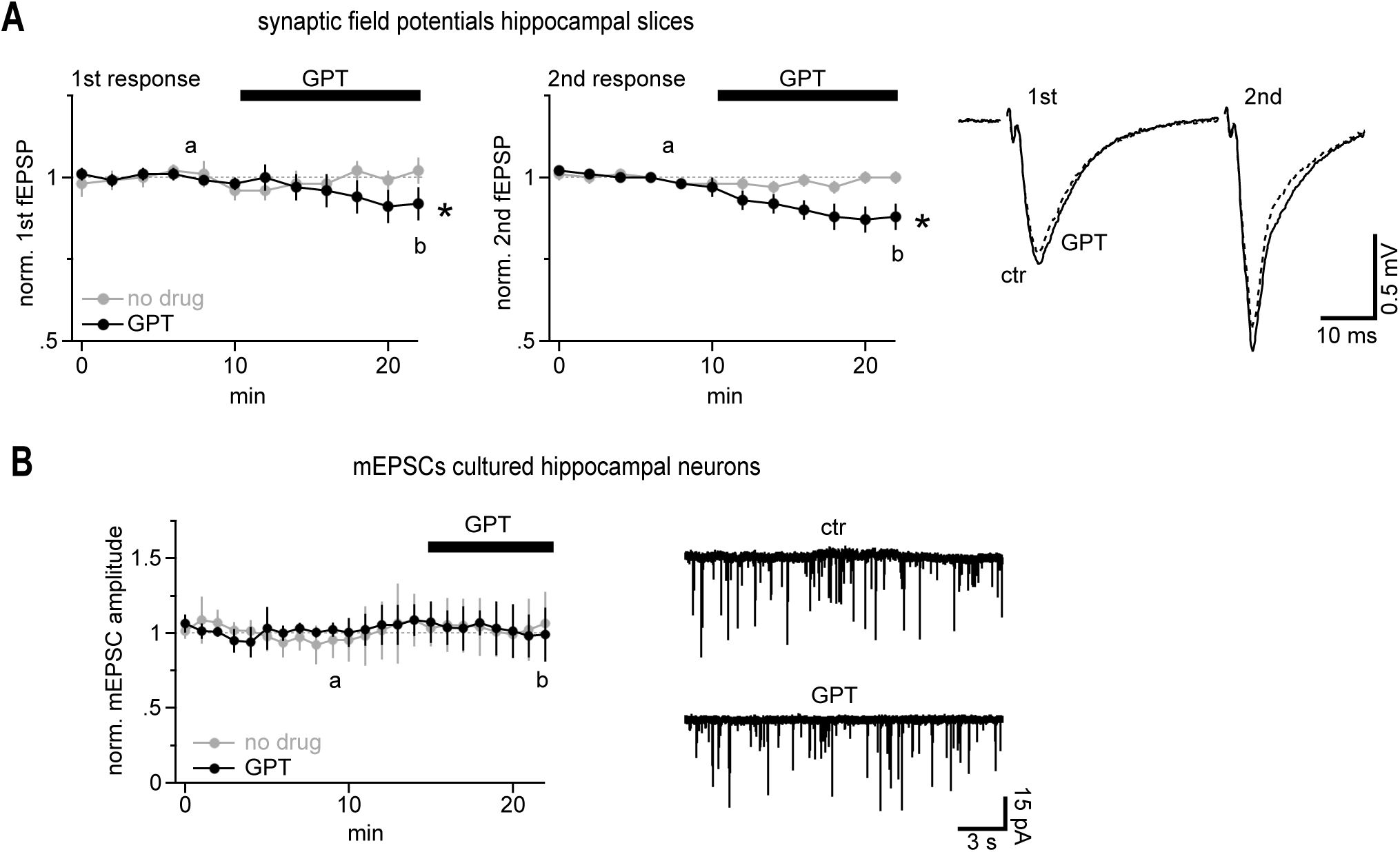
Synaptically released glutamate regularly co-activates neighboring synapses to a small extend. (A)AMPA receptor-mediated population synaptic responses in hippocampal slices are enhanced by glutamate acting on neighboring synapses. Left panel, summary of the 1^st^ slopes of fEPSPs recorded in CA1 Str. rad. Note the slight, but statistically significant decrease in fEPSPs upon application of the glutamate scavenger system (GPT, n=9; “no drug” control experiment with placebo solution exchange, n=11). Middle panel, the inhibitory effect of GPT is more pronounced on the 2^nd^, facilitated fEPSPs which accompanies a higher spatial density of releasing synapses. “a” and “b” denote the times of the example traces illustrated in the right panel. Right panel, example traces illustrating the effect of GPT on population synaptic responses. (B) The glutamate scavenger system GPT is too slow to inactivate glutamate immediately after release in the synaptic cleft; the amplitude of miniature EPSCs remains unaffected. Miniature EPSCs were recorded in dissociated cultured neurons. As the nearest neighbor distance of synapses in cultured neurons is too large (>= 1 µm) to allow for cross-talk, the amplitude of these currents is a measure of intra-synaptic AMPA receptor activation only. “a” and “b” denote the times of the example traces illustrated in the right panel.

## Discussion

Our study provides experimental measurements of the distance from an individual synapse at which glutamate can activate a glutamate binding protein such as the glutamate sensor iGluSnFr. We show that putative single and multi-quantal release from small hippocampal synapses activates iGluSnFr molecules in a neighborhood with a radius of ∼2 µm. This neighborhood is much larger than expected based on previous theoretical models of glutamate spread in the neuropil following synaptic release [11,15]. In fact, when we employed these models and added iGluSnFr molecules according to [38,61] a single vesicle is predicted to generate a local iGluSnFr response of less than 1% DF/F at a distance of 1500 nm (Suppl. Fig.4) whereas we experimentally determined an iGluSnFr response of ∼5.4% DF/F at 1500 nm (cf Fig. 2). This means that our experimental results exceed the theoretical predictions by a factor of ∼5. Responses at this distance are sufficiently far away not to be contaminated by light originating from the activated synapse, as evidenced by the spatial restriction of the red channel in Fig. 1. Further, the spatial gradients of the signals at that distance compare well to the optical resolution such that the local iGluSnFr response amplitudes are well resolved. Thus, our optical recordings suggest that glutamate after vesicular release may penetrate much further into the peri-synaptic tissue than previously reported and may also imply a larger synaptic cross-talk component. This view of an unexpectedly large spread of synaptic glutamate into the extracellular neighborhood is supported by our scavenger experiments which suggest that even AMPA-R mediated synaptic communication is carried by synaptic cross-talk to a small extent (Fig. 7).

We further employed 2P-glutamate uncaging to quantitatively determine distance-dependent activation of AMPA and NMDA receptors situated on dendritic spines. It is inherent to this approach that the uncaging laser spot is much larger than a synaptic cleft, the laser pulse releases more glutamate than contained in a single vesicle and the duration of the laser pulse (0.6 ms) is longer than it takes to empty a synaptic vesicle. Therefore, the question arises what can uncaging experiments tell us about synaptic cross-talk and how do these differences affect our conclusions? The differences between synaptically generated glutamate gradients occurring on and very near a spine and those induced by uncaging on or close to a spine are particularly large: vesicular glutamate is released faster (vesicle content liberated within ∼0.1-0.3 ms [62]) and is initially confined to the synaptic cleft. This difference can most clearly be seen when considering that uncaging at 0 nm (at the spine) equals the AMPA-R amplitude caused by a single vesicle (∼12 pA) while we in fact photo-release the amount of ∼5 vesicles. Thus, uncaging amplitudes close to the source are smaller than if the same amount of glutamate would be liberated in the synaptic cleft and for this reason the distance-dependent curves appear “flatter” than they really are. Therefore, λ values cannot be taken to describe the relative spatial decay of synaptic cross-talk responses. However, the situation is different if we compare the remote action of synaptic and uncaging sources of glutamate. Synaptically released glutamate escapes the synaptic cleft and spreads within the neuropil like a progressively enlarging cloud and reaches the target spine with some delay. Thus, even after fast and very local vesicular release, the wave of glutamate arriving at a remote target synapse will be slowed, broadened and diluted. It is instructive to compare the glutamate concentration profiles arriving at a remote synapse, eg at 500 nm, after brief vesicle release (<0.1 ms) and after uncaging release from a 3D point-spread function (PSF) for 0.6 ms. For this comparison we simulated the two types of release and the ensuing neuropil diffusion of 5000 glutamate molecules according to the standard approaches used by [11,15]. Suppl. Fig. 5A clearly shows that after vesicular release the glutamate concentration reached a ∼2-fold higher peak at 500 nm when compared to prolonged 0.6 ms release from a PSF volume (x/y FWHM = 280 nm, ω_z_ ∼ 3.5 ω_x,y_). Adding AMPA-receptors to the simulation at 500 nm (following [11,15]) demonstrated that synaptic release also causes 2-fold stronger glutamate receptor opening when compared to uncaging the same amount of glutamate (Suppl. Fig. 5B). Thus, uncaging glutamate at 500 nm significantly underestimates the AMPA-R-mediated synaptic cross-talk at the same distance.

During uncaging we release the glutamate content of ∼5 vesicles (∼35000 molecules) and at a distance of ∼500 nm this generated an AMPA-R-mediated uEPSC of ∼4 pA (cf Fig. 3 & 4). Thus, when keeping in mind that synaptic receptor opening will be even stronger, our data suggest that a 5-vesicle release event from a synapse at the average nearest neighbor distance of ∼500 nm can cause a cross-talk current of at least ∼4 pA which represents ∼33% of the single quantal response of synaptic AMPA-Rs (19 pA). Further, when glutamate transporters were blocked (tfb-TBOA, Fig. 4D) our uncaging data suggest a 5-vesicle release event to cause a cross-talk current of at least ∼6.4 pA (53% of quantal amplitude) at neighbor synapses located at 500 nm. This large cross-talk component of distant uncaging is not predicted by standard models of glutamate diffusion in the neuropil. To illustrate this, we followed the modeling approach of [15] which calculates a synaptic open probability (P_o_) of ∼0.172 for the release of single vesicle filled with 7000 molecules of glutamate (with and without transporters). We added a three dimensional, PSF-shaped and 0.6 ms-lasting source of 35000 molecules of glutamate (5 vesicles) and calculated the resulting open probability of AMPA-Rs at 500 nm (Suppl. Fig. 5C). It can be seen that this model predicts at this distance in response to uncaging a P_o_ of ∼0.022 (Suppl. Fig. 5C) in the absence of transporters) which represents only ∼13% of the synaptic response (0.172/0.022) whereas our uncaging responses reached 53% of the quantal amplitude (6.4 / 12 pA). Thus, our experimental data exceed the theoretical estimates by a factor of ∼4. Note that other models predict a much high P_o_ for synaptic AMPA-R (up to 0.7 [11]) which would make the difference between our experimental observation and the theoretical predictions even larger.

How relevant is a cross-talk signal occurring after multi-vesicular release for synaptic excitation of networks? While it was assumed for a long time that at most one vesicle would be released per action potential from small cortical synapses [63,64], there is now strong and growing evidence that many synapses, if not all, typically release up to 5 vesicles [39,43-46]. Multi-vesicular release from and cross-talk between Schaffer collateral synapses is consistent with our glutamate scavenger experiments in this pathway (Fig. 7) and suggests synaptic cross-talk may be more relevant for neuronal information processing than thought previously.

Does glutamate uncaging saturate glutamate uptake mechanisms? λ_AMPA_ did not decrease if we lowered the amount of glutamate released by uncaging (cf Fig. 6C) as would be expected if transporters were overwhelmed. Similarly, small synaptic iGluSnFr responses (selected small & spontaneous events) showed a glutamate spread comparable to that of large responses (Fig. 1&2). On the other hand, the λ_AMPA_ and λ_NMDA_ were slightly enhanced when we uncaged more glutamate (Fig. 6), indicating that a larger glutamate load cannot be handled with similar efficiency and that the uptake system operates close to the border of linearity when challenged by the standard uncaging pulse. This scenario explains why applying three uncaging spots simultaneously led to a supra-additive spread of glutamate (Fig. 5). The spread of glutamate likely is facilitated because glutamate transporters start to be overwhelmed under this high local load of glutamate.

Can synaptic activity generate such glutamate load? In our uncaging experiment the spine head integrated glutamate release equivalent to ∼15 vesicles (3 uncaging spots, each 5 vesicles) within a radius of ∼0.75 µm corresponding to a volume of ∼1.8µm^3. This volume of neuropil on average contains ∼3-4 synapses (2 synapses/µm^3) each of them being able to release up to 5 vesicles. Thus, if a handful of neighboring synapses are active together and undergo multi-vesicular release, the boost of synaptic current by ∼30%, as observed during the uncaging experiment, can indeed occur. The physiological boost might even be larger because synaptic activity can occur simultaneously while uncaging pules in our experiment were for technical reasons limited to a synchrony of ∼2 ms. Further, when keeping in mind, as argued above, that following uncaging lower glutamate concentrations are reached due to the psf-shaped source and the prolonged release time (0.6 ms), even fewer (<15) co-released synaptic vesicles may generate the same level of amplification as seen during uncaging. Such high density of active synapses is likely not achieved across a larger region typically recruited for experimentally stimulated compound synaptic responses and may explain the conclusion that transporters are not overwhelmed by synaptic activity in previous work [65].

What exactly is the role of astroglial glutamate uptake in limiting the action range of glutamate around a synapse, on the scale of less than 2 µm and below 2 ms? Blocking glutamate transport competitively by tfb-TBOA strongly increased λ_AMPA_ by ∼75% demonstrating that glutamate transporters are important for limiting the spread of glutamate. However, the intracellular translocation of glutamate by transporters of hippocampal astrocytes shows a high temperature sensitivity (Q10 ∼2.5, [52]) but λ_AMPA_ only modestly decreased by ∼15% when elevating the recording temperature from 20°C to 32° C. This indicates that intracellular translocation of glutamate does not greatly contribute to limiting λ_AMPA_. A potential explanation is that the translocation process itself is too slow, even at near body temperature, to efficiently remove glutamate molecules on the scale of 2 µm and below 2 ms. Rather, it appears likely that transporters limit λ_AMPA_ not by translocating glutamate but by binding glutamate and successfully competing with glutamate receptors for binding the ligand, as has been proposed for glutamate dynamics within the synaptic cleft [66]. Binding by transporters is rapid, precedes translocation and is competitively blocked by tfb-TBOA and therefore is consistent with our experimental observations. This scenario suggests that the number of transporter binding sites exposed to the extracellular space in the microenvironment is sufficiently high to reduce λ_AMPA_ even without translocation of glutamate (cf. Fig. 5 [15]) but also that those binding sites can be partially depleted if the local density of active synapses grows high (see above). Conversely, it can be concluded that if the local transporter density is slightly up- or down-regulated, this will result in a shorter or larger action range of glutamate. This connection puts astrocytes in an ideal position to tune synaptic cross-talk by strategic positioning of glutamate transporter molecules on their membranes, as recently proposed to happen after induction of LTP [67].

Why has cross-talk via AMPA-Rs at cortical synapses not been reported before? Compared to NMDA-Rs, AMPA-Rs may be sensitive only to cross-talk from a handful of synapses in their immediate microenvironment and therefore resulting cross-talk amplitudes are substantially smaller (∼4 pA from a neighboring 5-vesicle release event). This makes AMPA-R mediated cross-talk more challenging to experimentally dissect. Yet, there are numerous observations in the literature which could also be explained by cross-talk but which have been neglected or interpreted in a different context: minimal stimulation paradigms in slices typically also generate a fraction of synaptic current responses which are much smaller than the quantal amplitude (eg <2.5 pA) and sometimes occur slightly delayed [68]. At least some of those responses could represent cross-talk from neighboring, co-stimulated synapses. Further, it is well known that quantal synaptic responses (eg Sr^2+^-induced or mEPSCs) show a broad amplitude distribution ranging from ∼20 pA down to a few pA and the magnitude of baseline noise, eg [69]. There are established factors that contribute to this variability such as dendritic filtering and heterogeneity of pre-/postsynaptic properties but it is not clear whether these can entirely explain the occurrence of very small quantal responses. The amplitudes of these small responses fall well within the range of what could be predicted for cross-talk responses due to release of nearby synapses - even for very short lambdas. Slowing extracellular diffusion increases the efficiency of glutamate to activate remote receptors [11]. A previous study experimentally slowed diffusion of synaptic glutamate by applying dextrans and observed that besides the expected increase in NMDA-R mediated current, AMPA-R-mediated mEPSCs were also slightly increased [25]. This could be caused by dextrans acting in the cleft and improving the availability of glutamate for AMPA-Rs, as reasoned by the authors [25], but it is also conceivable that some of the mEPSCs were arising from nearby synapses via crosstalk and were potentiated because dextrans also enlarged the spatial action range of AMPA-Rs. A related ambiguity is encountered when interpreting trial-to-trial variation of glutamate levels in the synaptic cleft as detected by rapidly dissociating AMPA-R antagonists. Facilitated synaptic currents were shown to accompany higher glutamate concentrations in the synaptic cleft [43,70]. One interpretation is that synaptic facilitation in combination with multi-vesicular release results in a higher cleft glutamate concentration [43] while another is that synaptic facilitation leads to a higher spatial density of active synapses and thereby to a larger cross-talk component [70]. While in our eyes both factors will contribute to the phenomenon and multi-vesicular release itself enhances cross-talk, it is very difficult to unequivocally differentiate between the two possibilities as reasoned by [71], in particular if the source of glutamate cannot be located and diffusional distances are unknown. In summary, future studies are needed to address whether and to what extent the above-mentioned examples are mediated by cross-talk. Nevertheless, these examples raise the possibility that the phenomenon of synaptic cross-talk has been encountered but might not have been fully recognized as such.

Our finding of small but regular synaptic cross-talk violates modelling studies which predict that cross-talk at AMPA-Rs is negligible and at NMDA-Rs at least two-fold smaller than what we observe experimentally [11,15-18]. To some degree this difference could be explained by more recent estimates of certain biological parameters such as a higher glutamate content of synaptic vesicles (7000-8000 molecules) [56,57], a wider synaptic cleft (>=24 nm) [72-74] and a deeper understanding of glutamate transporter reaction schemes [75]. A further shortcoming of existing modelling studies may be the approaches used to approximate diffusible signaling on the nanoscale in the brain. The extracellular space is usually modeled as a porous medium by averaging over local anatomical details and this has been very successful in predicting the spread of molecules in the brain over larger distances (>10µm) [76]. However, this approach is not suited to describe initial diffusion on a scale of less than 1 µm [77,78]. During this initial distribution in the neuropil molecules temporarily behave as being on lower dimensional spaces consisting of tunnels (1D) and sheets (2D) formed by the cellular elements [72] and it is conceivable that transmitter accumulation at a distance of 500 nm, a nearby synapse, is larger than predicted by the porous medium approach. However, more detailed analysis of such nano-diffusion is required to explore whether deviations from the porous medium approach together with updated model parameters (see above) can quantitatively explain the differences between our experimental results and previous simulations.

Limitations of the study. Our study employs 2P-scanning approaches at the resolution limit and our comparison to synaptic release relies on optical parameters, such as ε and the uncaging excitation volume, which are difficult to determine precisely in situ. Further, most of our experiments were performed at room temperature (∼20°C) and cross-talk phenomena in previous studies disappeared when employing higher, near-body temperatures [22,58](and other studies). Those questions of accuracy and how they relate to our interpretations are discussed below.

We estimated the FWHM of the uncaging point-spread function to be close to the theoretical optimum, ∼280 nm, by bleaching spines in slices (Fig. 6D-F). This approach may underestimate the true FWHM if partial bleaching of spines is followed by such rapid fluorescence recovery that we do not fully resolve the initial degree of the induced bleaching. However, an almost optimal beam configuration was also suggested by an independent method not relying on the exact PSF geometry, by FCS (Fig. 6H), which yielded an uncaging volume of 0.2 fl and also suggests an almost optimal configuration. FCS measurements were performed in solution and it is conceivable that due to scattering/blurring the excitation volume in the slices is slightly larger. However, this will partially be compensated for by the fact that the FCS-based estimate of the excitation volume takes into account all fluorescence fluctuations and some of them are of little physiological relevance for uncaging experiments. This is true particularly for the so called “wings” of the point-spread function which are most remote to the focal point [55]. Glutamate molecules released in those wings will strongly be diluted before they reach AMPA-Rs on the spine head and in so far, the FCS-based excitation volume estimate of 0.2 fl slightly overestimates the number of glutamate molecules contributing to the uEPSC (in the simulation shown in Suppl. Fig. 4 we assumed that all glutamate molecules, including those originating in the wings, are released within a 3D gaussian).

We estimated ε to be ∼0.3 as we noted that by only slightly elevating laser power, we can easily increase AMPA-R mediated uEPSC currents (cf Fig. 6C) up to ∼40 pA (not shown). If ε was significantly larger than 0.3, such strong amplitude increases are difficult to explain as we bathed the slices only in 5 mM MNI-glutamate (K_d_ AMPA-R ∼500 µM). On the other hand, the rapid rise of the uncaging EPSC suggested that at least several hundred µM glutamate acted at AMPA-Rs on the spine head. If we assumed a substantially lower fraction, eg ε ∼ 0.1, much slower rise times would be expected (eg [15], Fig.7), which would not be consistent with our observations. If ε was larger than 0.3, then our estimate of the number of uncaged glutamate molecules would increase proportionally. For example, if ε was 0.6, one uncaging pulse would correspond to ∼10 and not ∼5 vesicles. This would however not strongly affect our main conclusion as there is a safety factor of ∼2 when comparing uncaging to synaptic responses: AMPA-R opening following 0.6 ms uncaging is approximately two-fold less efficient than following the same amount of glutamate released in a synaptic fashion (Suppl fig 5).

Most of our experiments were performed at room temperature and a number of studies did not detect synaptic spill-over at near-body temperature [22,58]. We also assessed the effect of temperature on the spatial decay of uncaging EPSCs (Fig. 4E) and concluded that the effect of increasing temperature was relatively small. This contrasts the two previous studies [22,58] but this difference could be expected. Those studied looked at the activation of high affinity receptors (NMDA, mGluR) and the involved diffusion distances were - albeit not known precisely - likely much larger than ours induced by uncaging at submicrometer distances from the recorded spine. Glutamate transporters are much more effective in eliminating glutamate on longer diffusional distances [15](Fig 6 therein) which could explain the difference to our study.

On the whole, our results may challenge the view of exclusive point-to-point communication between neurons in the CNS by showing that synapses with multi-vesicular release rather regularly communicate or whisper to a cluster of ∼2-4 (AMPA-Rs) and up to >70 (NMDA-Rs) adjacent synapses and that this contributes to network activation as evidenced by our fEPSP recordings. Nearby synapses appear functionally coupled at a strength dropping with their distance. Functional coupling of closely spaced, but un-connected, neurons by cross-talk is a form of redundancy which decreases the theoretical storage capacity of the brain [4]. However, sacrificing storage capacity in favor of redundancy might enhance robustness: in case of irreversible damage of a postsynaptic neuron, information conveyed by incoming synapses would be lost without this redundancy, but cross-talk ensures that this information is still available to the network such that other, co-activated neurons can take over and re-route the information previously propagated by the lost neuron. Redundancy due to synaptic cross-talk could also be key to the well-known and remarkable stability and functional resiliency of the continuously learning brain: lifelong-learning artificial neural networks strongly suffer from so-called ‘catastrophic forgetting’ caused by the loss of learned and sparsely represented information being overwritten during new learning cycles [79]. Interestingly, the inclusion of a simulation of local neurotransmitter diffusion into the design of artificial neuronal networks helps to eliminate this unwanted phenomenon [79]and introduces some of the resilience known from the biological brain.

A further major aspect of our work is the finding of supra-additive spread of glutamate caused by coincident activity of nearby synapses. This finding suggests a new mechanism by which astrocytes can regulate synaptic integration on the millisecond time scale through acting on the extracellular space. While intracellular calcium signaling or gliotransmitter release by astrocytes is very slow and happens within seconds, astrocytes could tune the local volume density of glutamate transporters[80] and by this regulate high frequency neuronal activity: the density of transporters will set the degree and regionality of supra-additivity of kHz coincident neuronal activity and foster pseudo-clustered activity along the same and across different dendrites – a mechanism so far only known from dendritic spatio-temporal summation.

Taken together, our results suggest that a deep functional understanding of neuronal circuits and behaviors not only calls for deciphering synaptically connected pairs of neurons in the brain but may also require considering the immediate spatial neighborhood of neurons and their synapses on the sub-micrometer scale.

## Acknowledgments

We are grateful for the assistance of the Viral Core Facility of the Medical Faculty of the University of Bonn, supported in part by SFB1089.

## Materials & Methods

### Animals

All procedures were planned and performed in accordance with the guidelines of the University of Bonn Medical Centre Animal-Care-Committee as well as the guidelines approved by the European Directive (2010/63/EU) on the protection of animals used for experimental purposes. All efforts were made to minimize pain and suffering and to reduce the number of animals used, according to the ARRIVE guidelines. Mice were housed under a 12h light–dark-cycle (light-cycle 7am/7pm), in a temperature (22±2°C) and humidity (55±10%) controlled environment with food/water *ad libitum* and nesting material (nestlets, Ancare, USA). Animals were allowed at least 1 week of acclimatization to the animal facility before surgery and singly housed after surgery. Male and female C57Bl6/N mice (Charles River, Sulzfeld, Germany) were used between the ages of P15 and P20, except where other ages are noted in the results.

### Slice preparation

Animals were anesthetized with Isofluorane gas, decapitated, and the brain was removed and submerged into ice cold dissection solution (in mM): 87 NaCl, 2.5 KCl, 1.25 NaH_2_PO_4_, 7 MgCl_2_, 0.5 CaCl_2_, 25 NaHCO_3_, 25 glucose, 75 sucrose (gassed with 95%O_2_ /5% CO_2_). Frontal or ventral horizontal slices (300 µm thick) were made on a vibratome (Leica VT 1200 or Thermo Scientific HM650V) and incubated at 35°C for 30 minutes in a submerged chamber filled with the dissection solution. Slices were then transfered to a holding chamber filled with oxygenated artificial cerebral spinal fluid (ASCF) until the experiments began. The ACSF contained (in mM) : 124 NaCl, 3 KCl, 1.25 NaH_2_PO_4_, 2 MgCl_2_, 2 CaCl_2_, 26 NAHCO_3_, and 10 Glucose, pH 7.4 (Sigma-Aldrich), and was continuously bubbled with 95%O_2_ /5% CO_2_. This solution was used for perfusion during the subsequent electrophysiology recording and imaging experiments.

### Electrophysiological recording

Slices were positioned in a recording chamber on the stage of a microscope and perfused with the recording solution which routinely contained 25µM APV and 10µM TTX or other blocker cocktails as stated in the text. Patch pipettes were pulled on a vertical puller (Narashige PP-830) with a resistance of 4.5-6 MΩ. Pipette solution for the voltage clamp experiments contained (in mM): 125 K-gluconate, 4 Na_2_-ATP, 2 MgCl_2_, 10 HEPES, 20 KCl, 3 NaCl, 0.5 EGTA (pH = 7.3, 280-290 mOsm) and 25µM Alexa 594 or 100 µM Tetramethyl-rhodamine (TMR, for experiments in Figure 3 we used 400 µM TMR) to visualize spines. For experiments to determine the spatial range for NMDA receptors, Cs-based pipette solution was used. It contained (in mM): 130 CsOH, 15 CsCl, 130 D-Gluconic acid, 2 MgCl_2_, 10 HEPES, 0.5 EGTA, 5 QX314 (pH = 7.3, 280-290 mOsm) and 25 µM Alexa 594. Holding potential was set at −65 mV, except for experiments using Cs-based pipette solution, which was set at +40 mV to unblock NMDA receptors from Mg_2+_. Electrophysiological data for quantifying the spatial range of uncaged glutamate via glutamate uncaging was acquired on a 2-photon rig equipped with an Ultima multiphoton microscope (Bruker) with two independent pairs of scanning mirrors coupled to two Chameleon vision II lasers (Coherent). The amplifier was an EPC-10 (HEKA), controlled by PatchMaster software (HEKA), triggered by the PrairieView software (Bruker) to coordinate uncaging and imaging lasers, as well as the recording software. The electrophysiological data was sampled at 20 kHz and filtered at 3 kHz. Imaging data for the optical reporter of synaptically released glutamate (iGluSnFr) was acquired on a Nikon A1R MP 2-photon scanning microscope (Nikon) equipped with a BVC-700 (Dagan) amplifier and using WinWCP software (Strathclyde) for current clamp recording. The mEPSC response to enzymatic scavengers of glutamate were recorded on a conventional electrophysiology rig equipped with an EPC-7 amplifier (HEKA) using pCLAMP 9 software (Molecular Devices). fEPSP data was acquired as follows: Animals were anesthetized with Isoflurane (Baxter) and decapitated. The brain was removed from the skull and chilled for 1 min in cooled (4°C) artificial cerebrospinal fluid (ACSF) containing in mM: 125 NaCl; 2.6 KCl; 1.4 MgSO_4_; 2.5 CaCl_2_; 1.1 NaH_2_PO_4_; 27.5 NaHCO_3_ and 11.1 D-glucose; pH 7.3, 310 mosm/kg. The hippocampus was transversally cut in 400 μm slices (VT1200S, Leica). Slices were equilibrated in a custom-made submerged chamber in ACSF continuously gassed with carbogen (95% O_2_; 5% CO_2_) for 30 min at 32°C, and subsequently kept at RT. fEPSP recordings were obtained from P15 – P25 animals. During experiments, slices were continuously super fused with ACSF supplemented with 10 mM HEPES and 2 mM sodium pyruvate. Glutamate-pyruvate transaminase was applied in a concentration of 5 U/ml. Paired fEPSPs with an interstimulus interval of 40 ms were evoked by stimulating Schaffer collaterals at 0.033 Hz with a pulse duration of 0.2 ms. fEPSPs were recorded in the stratum radiatum of the CA1 region using glass microelectrodes (Science Products, Hofheim, Germany) filled with ACSF. Data was acquired using a Multiclamp 700B amplifier (Axon Instruments), digitized on a Digidata 1322A (Axon Instruments) and stored on a PC. All experiments were performed at room temperature (22–24°C). fEPSP slopes were used as a measure of dendritic activity and determined between 20-80% of the maximum field amplitude.

### Glutamate uncaging

To ensure accurate and repeatable results, the following alignment and calibration steps were performed every day. After the lasers were turned on and warmed up, the beam alignment for the imaging and uncaging lasers was checked at the objective using a fluorescent target. Laser power was controlled via a Conoptics EOM and was measured at the objective with a slide power meter (Thorlabs) to generate an uncaging power calibration curve for the day. The correspondence between the x-y pointing for both sets of scan mirrors was calibrated using a fluorescent plastic slide, and any required adjustments were entered in the PrairieView control software. Uncaging laser wavelength was 720nm; imaging wavelength was determined by the experiment requirements as stated in the main text. MNI-Glu (Tocris) was prepared by dissolution into ACSF at 30 mM, then aliquoted in 100uL portions and frozen. Aliquots were thawed immediately before addition to the recording chamber, and were never recycled/reused after thawing.

Cells were voltage clamped in the whole cell mode for 10 minutes to allow filling of dendritic spines. Only spines and dendritic shaft segments at a depth of 20-30µm from the tissue surface were targeted to ensure that the uncaging power was not attenuated differently by scattering and other effects between experiments. Once a dendrite at the correct depth and orientation was found, perfusion of the recording solution was halted, and 300µL of ACSF containing 30mM 4-Methoxy-7-nitroindolinyl-caged-L-glutamate (MNI-caged glutamate, caged glutamate) was carefully pipetted into the recording chamber. The bath volume before application of caged-glutamate was kept at 1.5mL, such that after addition of caged-glutamate the final concentration of MNI-Glu in the recording chamber was 5mM. The recording experiment resumed after the caged compound had been in the bath for 10 minutes to allow complete diffusion into the slice. A high resolution z-stack of the targeted spine was taken to ensure that there were no dendrites or spines above or below the targeted spine from the same cell. Uncaging points were positioned orthogonal to the broadest part of the spine head and parent dendrite using the imaging laser, and the position was rechecked automatically following each uncaging protocol to eliminate any experiments where movement artefacts may have influenced the relative distance between the uncaging point and the spine head. Typical experiments used 10 uncaging points at 1Hz, spaced 100nm apart, and beginning uncaging at the furthest point from the spine head. Uncaging pulses were 0.6ms in duration; the power was set so as to elicit an uEPSC at the spine head of nominally 12 pA (accepting 10-14pA due to trial-to-trial fluctuations). This amplitude matches the size of mEPSCs commonly reported and also observed in the lab (data not shown). The laser power value required for this amounted to ∼23 mW and was used thrughout the study.

### Fitting uEPSCs

To optimally measure the amplitude of even small uEPSCs, the recorded currents were fitted with a difference of two exponential functions being defined by two time constants describing rise (2.3 ± 0.3 ms) and decay times (9.1 ± 0.6 ms, n=27) and a scale factor describing the amplitude.

### Induction of SE by suprahippocampal kainic acid application

Surgery, induction of SE and postoperative care were previously described in detail {Pitsch, 2019 #2369}. Briefly, 70 nl kainic acid (20mM, Tocris) was injected above the left hippocampal CA1 region (−2AP −1.4ML −1.1DV, {Paxinos, 2012 #2224}) of anaesthetized [16 mg/kg xylazine (Xylariem, ecuphar) and 100 mg/kg ketamine, i.p. (Ketamin 10%, WDT)] 15 adult male C57Bl6/N mice. Control injections were performed with the same volume of 0.9% NaCl. Animals were used 5-9 days following the injection.

### Determination of optical resolution/width of the point-spread-function (psf) of the uncaging laser spot

A common test to estimate the width of a psf is to image/scan a sub-resolution sized fluorescent bead such that the apparent size of the bead in the image reflects the dimension of the underlying psf. A fit of a gaussian function to the diameter of the scanned bead can be used to extract and calculate the full width at half-maximum fluorescent intensity (FWHM). Here we aimed to determine the resolution of our optical system in situ, near a spine in the slice, and used a dye-filled spine itself to probe the shape of the psf. The level of laser-induced bleaching of the spine was used as a read-out of the overlap of the psf of the uncaging laser with the spine. The level of bleaching will rise the closer we bring the uncaging/bleaching spot to the spine and will reach a maximum (spine>psf) when the psf is fully contained in the spine (in xy plane, see Fig. 6F). Therefore, if a gaussian is taken to estimate the shape of the psf, the increase in level of bleaching when bringing the psf/laser spot closer to the spine will follow the integral of that part of the gaussian which overlaps and bleaches the spine. The level of bleaching (“norm. bleaching”, Fig. 1) was fitted with a cumulative distribution function of a gaussian yielding the FWHM of the underlying gaussian describing the psf:

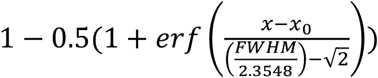

### Fluorescence correlation spectroscopy to determine the excitation volume of the optical uncaging system

We used two-photon fluorescence correlation spectroscopy recordings acquired with the Ultima multiphoton microscope (Bruker Corporation, Billerica, USA) as employed for uncaging experiments (720 nm, 60x Nikon NIR Apo water objective NA 1.0) to estimate the uncaging excitation volume. The laser power at the sample was measured and kept between ∼ 5-7 mW. Emitted fluorescence was filtered by an IR-blocker and a bandpass filter 550/100 (AHF analysentechnik AG, Tübingen, Germany) before being detected by a cooled PMT (bh HPM-100-40 Hybrid Detector, Becker & Hickl GmbH, Berlin Germany). Three recordings (each 120 s) were performed in 1.5 ml of a 50 nM tetramethylrhodamine-dextran (D3307, Thermo Scientific, Waltham, USA) dissolved in ddH2O at 18 °C while parking the laser beam in the center of the scan field. Time-correlated single photon counting was performed using the bh SPC-150 module and bh SPCM software (version 9.66, Becker & Hickl GmbH, Berlin, Germany). Autocorrelograms were calculated from photon arrivals times and fitted using the FFS data processor 2.7 (SSTC, Minsk, Belarusian state university) using the standard 3D-diffusion model:

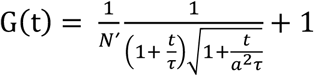

in which N’ is the apparent average number of molecules, τ is the translational diffusion time, *t* is the lag time and 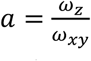, *ω*_*xy*_ and *ω*_*z*_ being the lateral and axial 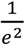 widths of the 2P point-spread function, respectively (*a* was set to 3.43). Residuals of the fit as shown in figure 6 were calculated by dividing the difference between the autocorrelogram and its fit by the standard deviation of the autocorrelogram. The standard deviation was computed based on dividing the fluorescent intensity trace in appropriate subsets according to ^1^. N’ provides an estimate of the number of fluorophores in the effective detection volume. As this the effective detection volume is an open volume without physical walls, fluorophores are moving across this boundary and contribute to the number of collected photons even when they have left the actual geometrically defined volume of the point-spread function ^2^. Therefore V_eff_ > V_psf_ and also for the number of fluorophores in V_psf_, N: N’ > N. N can be calculated from N’ by multiplying with the γ-factor, γ = 1/√8, a geometric factor that depends on the spatial shape of the detection profile ^2,3^. In our case (N ∼ 16, 50 nM TMR-Dx3kD) V_psf_ = V_eff_ * γ = 0.5 fl * 0.354 ∼0.2 fl.

### iGluSnFr detection of glutamate diffusion

The optical glutamate sensor iGluSnFr was expressed by using a mix of AAV1 and AAV5 viral vectors under control of the synapsin promoter (hSyn.iGluSnFr.WPRE.SV40; Penn State Viral Vector Core). Anesthetized [16 mg/kg xylazine (Xylariem, ecuphar) and 100 mg/kg ketamine, i.p. (Ketamin 10%, WDT)] juvenile male C57Bl6/N (Charles River Laboratories) mice (5-7 weeks old) were stereotaxically injected bilaterally into both ventral CA3 hippocampal regions (stereotaxic coordinates relative to Bregma: −2.5 AP, ±3.0 ML, −3.0 AP; 1 µL of undiluted virus; appr. titer: 8-10 × 10^12^) as described previously by using a beveled needle nanosyringe (nanofil 34G BVLD, WPI) under control of a micro injection pump (100 nl/min, WPI) ^4^.

Brain slices with strong expression in the Str. radiatum of CA3 and the hilus were selected for the experiment. To measure the spatial range of glutamate diffusion, a scan line (940nm) was placed in the Str. radiatum either parallel or perpendicular to the primary dendrites of CA3 cells, with a length of 10-15µm. A single uncaging spot (wavelength 720nm) was placed in the middle of the scan line and the uncaging laser pulse was triggered after a baseline of 200 lines was captured (∼1ms per line). For these experiments, the uncaging power was set to 20-25mW at the objective. The affinity of iGluSnfr is reported to be 4.9 µM, which is comparable to the affinity of NMDAR.

For measuring the spread of synaptically released glutamate from mossy fiber boutons, slices were similarly prepared and selected. These experiments were conducted on the Nikon A1 R two-photon system with only a single imaging laser, imaging wavelength was 920nm to excite both the red morphological dye and the iGluSnfr. Dentate granule cells were patch clamped in current clamp configuration with internal solution containing 400µM TMR. After holding the cell for 10-15 minutes, the axon would begin to fill with red dye, and complete, uncut axons were traced into the hilus region where iGluSnfr expression was strongest. The bath solution contained CNQX (10µM), 4-AP (100µM), and DCPCPX (1µM). A line scan was positioned crossing a clearly labeled presynaptic bouton, and somatic current injections of 1nA, 0.5ms were used to elicit action potentials.

### Estimation of spatial range with PSD95-gCamp5

The plasmid for the optical Ca^2+^ sensor GCamp6f fused to PSD95, pLenti-PSD95-GCamp6f. Lentiviral particles were prepared as previously reported (van Loo et al., 2019). Viral injections were performed as described above by using stereotaxic coordinates −1.9 AP, ±1.5 ML, −1.5 DV to target the dorsal hippocampal CA1 region. Imaging experiments were performed 2 weeks following virus injection.

Brain slices with strong expression in the CA1 Str. radiatum were selected for the experiment. Frame scans (excitation wavelength 950 nm) of the Ca^2+^ sensitive PSD95-Gcamp6f fluorescence was acquired at ∼40 ms, 110 nm spatiotemporal resolution. Glutamate uncaging was performed as previously described in the presence of 15 μM glycine to allow activation of NMDA receptors at negative potentials. Regions of interest were selected at a depth of ∼25 μm below the surface. After a 600 ms baseline acquisition, a single uncaging pulse was delivered in the center of the field of view. Responding spines could be detected as an increase in the local ΔF/F_max_. Fluorescence intensity, spatially averaged over manually selected spines (n=50 spines, 10 ROIs), increased rapidly in response to an uncaging event (maximal response at 80-120 ms after uncaging) and returned slowly to baseline. For many spines, resting fluorescence is undetectable above background. This precludes counting the total number of spines and calculating the fraction of activated spines in a field of view. Therefore, we conducted a pixel-based analysis to quantify the distance from uncaging site-dependence increase of Gcamp6f fluorescence: The ratio of activated pixels (ΔF greater than 4 SD in response to glutamate uncaging) at a certain distance over the total number of pixels at that distance (in the scan) was plotted versus distance from the uncaging site and used as an alternate metric for measuring the action range of uncaged glutamate at NMDA receptors.

### Simulation of Glutamate cross-talk

The effect of glutamate cross talk within a population of physiologically realistically distributed synapses was estimated by Monte Carlo simulation. Simulated synaptic elements consisted of a 112 nm radius “hard-core” postsynaptic density with a dedicated presynaptic partner. All synapses were assumed to be identical with respect to release probably and quantal response amplitude (12 pA). One quantal equivalent was defined as the response of a single postsynaptic element to release from its dedicated presynaptic element without any cross-talk. Spherical volumes with 15 μm radius were sequentially populated with synaptic elements in a random manner. For this candidate synaptic elements were generated with geometric centers sampled from a uniform random distribution, checked against all other synaptic elements in the simulation volume and placed into the simulation volume if it would not collide with an existent element. Synaptic elements were added in this manner until the synaptic density reached 2.07/μm^3^. This approach was previously used by ^5^. The amplitude of glutamate cross-talk/spillover responses between synaptic elements was modeled as exponentially decaying with inter-synaptic distance. A subpopulation of active pre-synapses was randomly selected from the entire population. To avoid edge effects, the responses were evaluated only at synaptic elements at least 10x spatial range from the edge of the simulation volume. For a spatial range of 450 nm, a representative simulation of a 15 μm radius volume contained 19972 spines greater than 4.5 μm from the edge of the simulation volume. 60 independently generated volumes were simulated, providing simulated responses for ∼1.2 million spines.

### Analytical description of spatial summation of activation at a linearly responding, non-saturable spine

We may also consider the analytical solution to how cross-talk builds up at an individual spine. Let Q be the quantal response at a “hard wired” synapse, λ spatial action range of glutamate, and n the density of synapses. Then, the infinitesimal activation increment of a spine at the origin due to all glutamate release happening (from all synapses) in the volume element dv at r’ is given by:

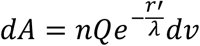

The summed activation of the spine due to concerted network activity is given by the volume integral:

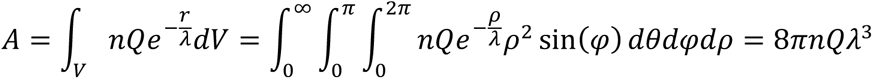

### Isotropic spread of iontophoretically injected glutamate in the neuropil of CA1 Stratum radiatum

The glutamate sensor iGluSnFR was virally expressed in astrocytes (AAV1.GFAP.iGluSnFr.WPRE.SV40, Penn State Viral Vector Core). Anesthetized [100 mg/kg ketamine (Ketamin 10%, betapharm) + 0.25 mg/kg medotomidine (Cepetor, CPPharma) i.p.] C57Bl6/N (Charles River Laboratories) mice (4 weeks old) were stereotaxically injected bilaterally into both ventral hippocampi (stereotaxic coordinates relative to Bregma: −3.5 AP, ±3.0 ML, −2.5 AP; 1 µL of undiluted virus) as described above. Finally, anesthesia was stopped by i.p. injection of 2.5 mg/kg atipamezol (Antisedan, Ventoquinol). To ensure analgesia, 5 mg/kg carprofen s.c. (Rimadyl, Zoetis) was injected for 3 consecutive daysAcute hippocampal slices (300 µm thick) were prepared after two to four weeks after virus injection. Experiments were performed in the presence of the glutamate receptor inhibitors NBQX (20 µM), D-APV (50 µM) and LY341495 (100 µM) and the sodium channel blocker TTX (1 µM) at a temperature of 34 °C. 2P excitation fluorescence microscopy of was performed as described previously (Anders et al. 2014). An iGluSnFR-expressing astrocyte in the CA1 S*tratum radiatum* and a region of interest for line scanning in the periphery of the cell were pseudo-randomly chosen. Line scanning of iGluSnFR fluorescence was performed as illustrated in Suppl. Fig. 2 at a frequency of 300-500 Hz and glutamate was applied iontophoretically (npi, Germany) close to the middle of the scanned line (∼1 µm) for 250 ms. In each experiment, line scans were performed both in parallel and perpendicular to the CA1 pyramidal cell layer. The iontophoretic current was 10 nA. We verified in each experiment that much larger iontophoretic glutamate injections (∼100 nA) were needed to saturate iGluSnFR. The background fluorescence was subtracted from line scan data. The latter was processed and analysed as illustrated and described in Suppl. Fig. 2 and its legend.

---

**Suppl. Figure 1:**
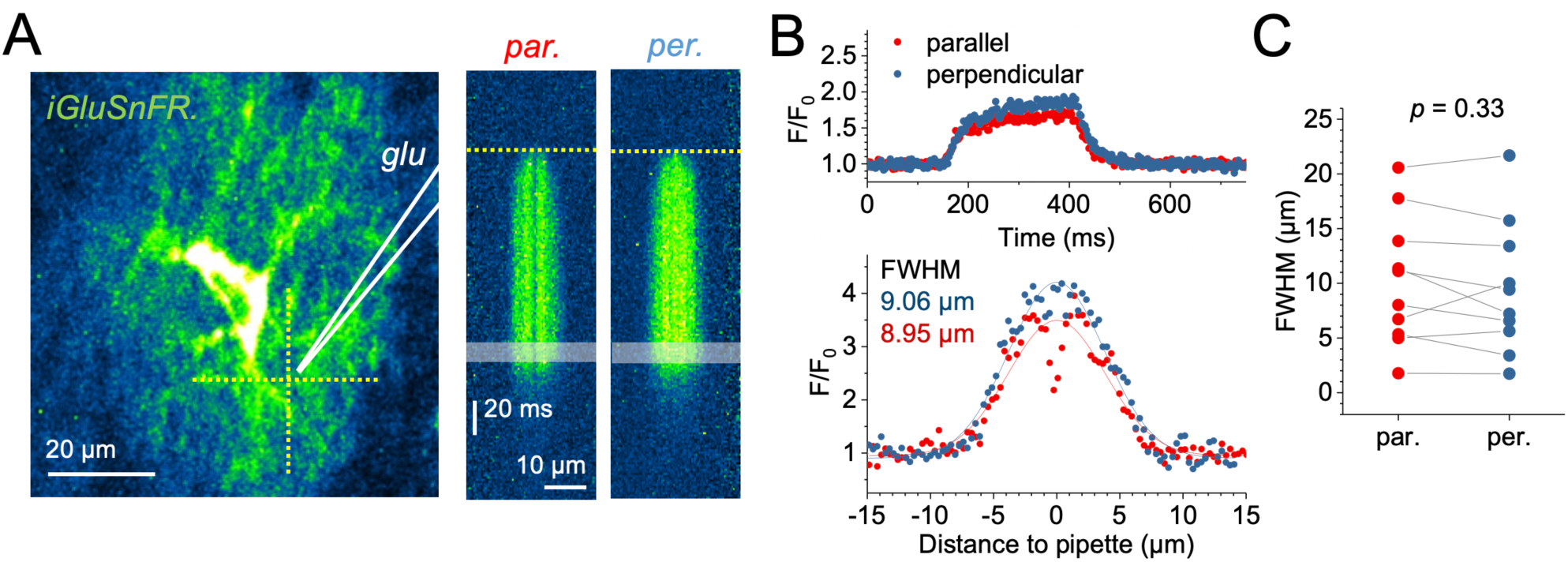
Isotropic spread of glutamate visualized by monitoring of iGluSnFr expressed by astrocytes. Long iontophoretic glutamate applications were used to minimize the potential buffering effect of membrane-anchored iGluSnfr molecules. (A)Schematic of the experiment (left panel, glutamate iontophoresis pipette in white, iGluSnFR expressing astrocyte in green). Line scans of iGluSnFR fluorescence were obtained in parallel and perpendicular to the CA1 pyramidal layer (left panel, dotted yellow lines) and glutamate was applied for 250 ms (current 10 nA). For analysis and display, the baseline fluorescence intensity (F_0_) was determined in a 100 ms time window for each x-coordinate (lane) and the ratio F/F_0_ was calculated for each lane (middle and right panel, start of iontophoresis illustrated by yellow dotted lines). iGluSnFR saturation was not observed in these experiments and usually required a much stronger glutamate injection (∼100 nA, verified in each experiment). (B) Example of iGluSnFR fluorescence over time during glutamate application (top panel, averaged over entire line scan). The spatial profile of iGluSnFR fluorescence during the last 20 ms of iontophoresis (shaded area in line scans in **A**) was analysed to estimate the spatial spread of glutamate in both directions. Spatial spread was quantified by the full width at half maximum of a Gaussian distribution fitted to the fluorescence profiles (lower panel). (C) No statistically significant difference between both directions were observed (n = 10 recordings in 10 different hippocampal slices, parallel and perpendicular line scans always paired, paired Student’s t-test).

**Suppl. Figure 2:**
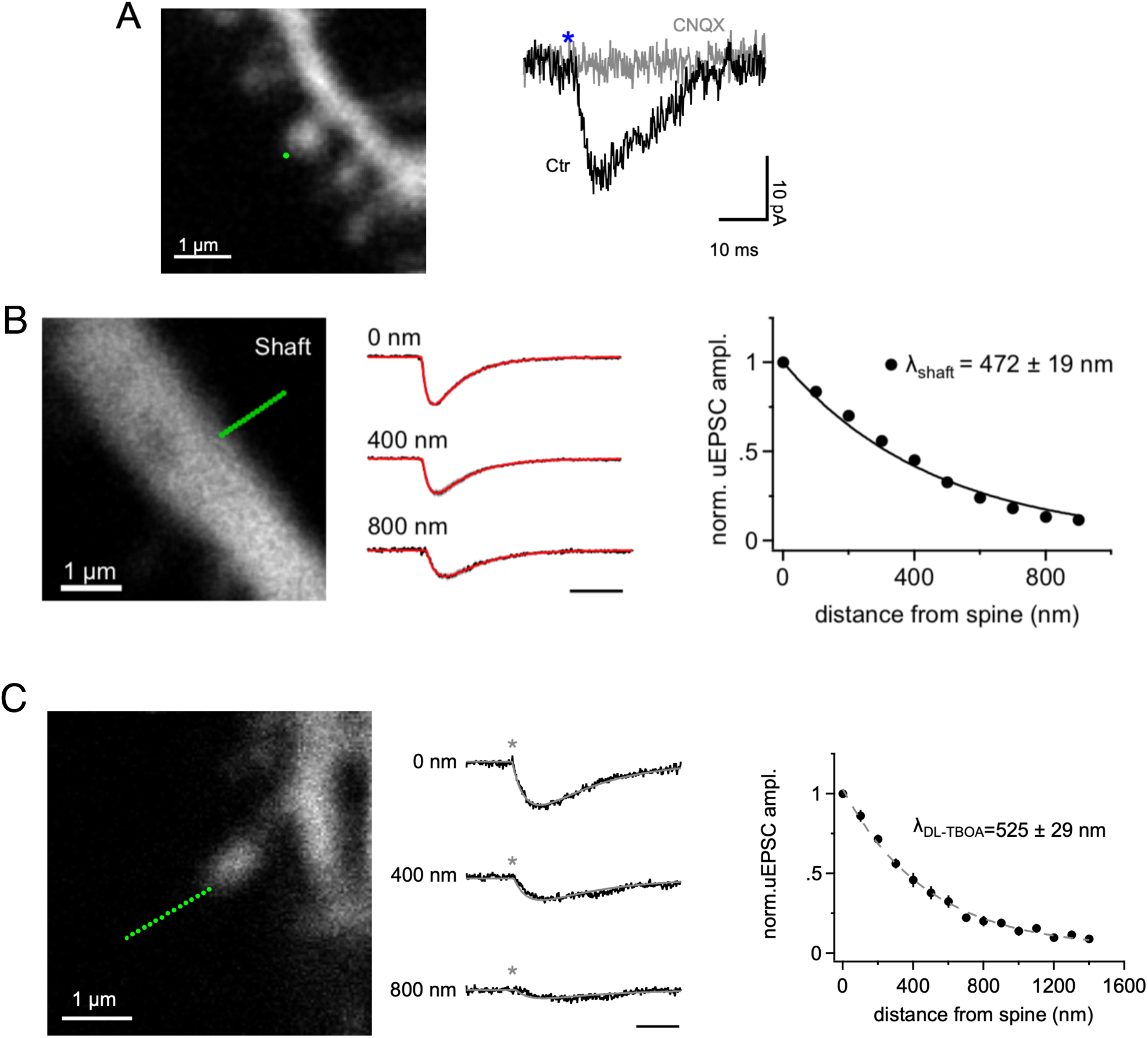
uEPSCs are mediated by AMPA receptors, the SARGe is similar at dendritic shafts and weaker blockade of transporters shows a mild increase in SARGe. (A)Uncaging EPSCs (uEPSCs) were mediated through AMPA receptors. A single uncaging spot placed on the edge of the spine head (green dot, upper panel) while NMDA receptors were blocked by 25 µM APV in the bath, elicited an AMPA-mediated uEPSC similar to miniature EPSCs (black trace, lower panel). The blue asterisk indicates the time of the uncaging pulse. After applying 20 µM CNQX in the bath, the same uncaging pulse did not elicit any response at the same spine (grey trace, lower panel, n = 5). (B) SARGe at dendritic shaft-located AMPA receptors, λ_shaft_, equals the SARGe measured at spine heads, suggesting the SARGe is determined by the surrounding neuropil and not by the type or location of the postsynaptic site. The precision of positioning the uncaging laser spot and the accuracy of the mutual alignment of both scanning systems (imaging and uncaging) was experimentally verified (Suppl. Fig. 3), scale bar, 10 ms, responses peak normalized. (C) Glutamate uncaging in the presence of the weaker glutamate transporter blocker DL-TBOA (100 µM, n=13). Partial blockade of glutamate transporters increases the SARGe showing that λ_AMPA_ also is sensitive to sub-maximal reductions in the number of unoccupied transporters. Scale bar 10 ms.

**Suppl. Figure 3:**
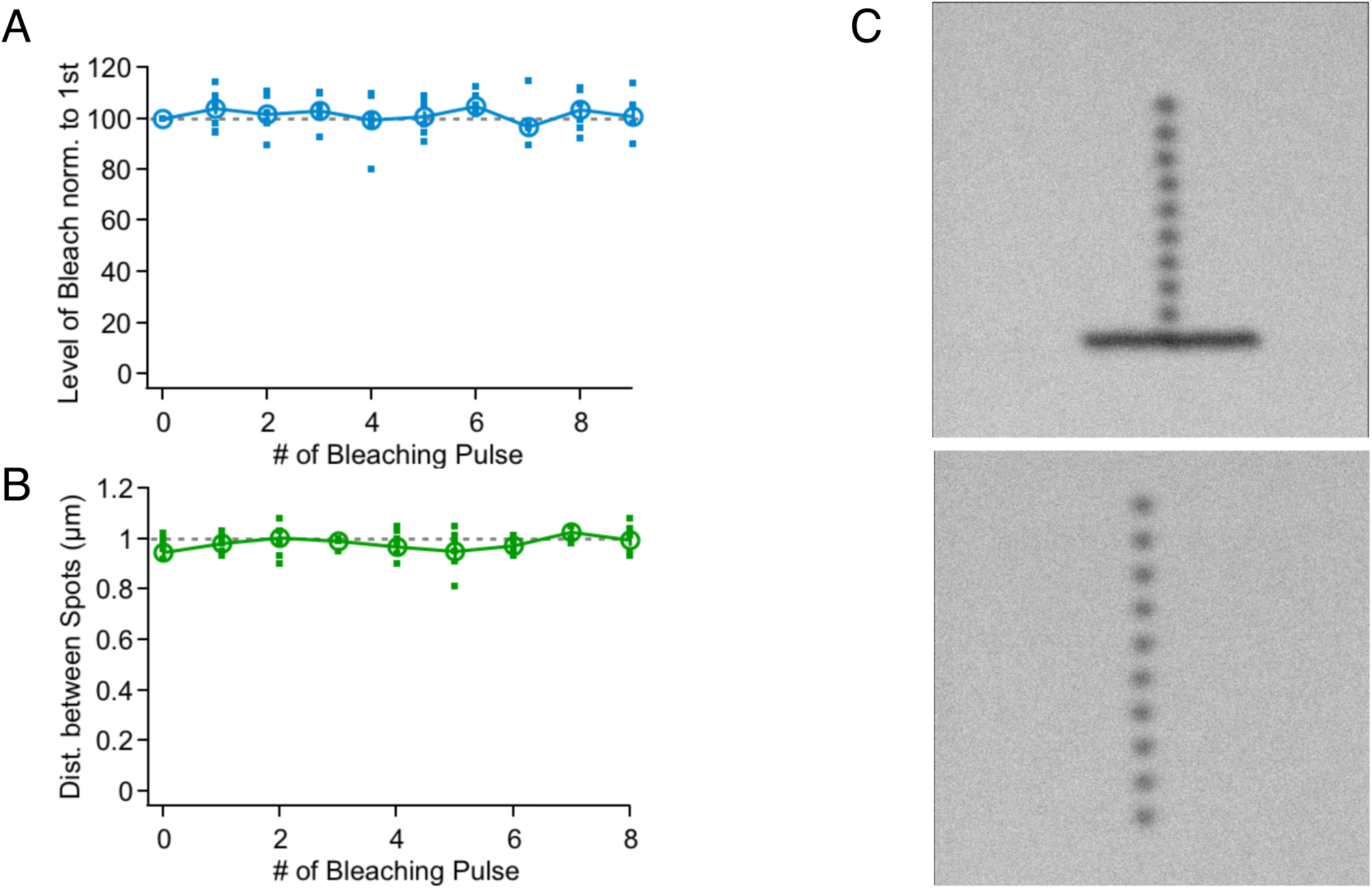
Verifying reproducibility and precision of applied uncaging power and spot positioning. (A)A plastic slide (Chroma Corp.) was used to visualize programmed uncaging events as bleached spots (cf C)). The level of bleaching as a readout of uncaging laser power was quantified in a series of uncaging spots and were constant with very little variability. (B) Uncaging spots were positioned at nominally 1 µm intervals (C) lower panel) and image analyses of a scan of a bleached plastic spot revealed the high precision of spot placement. (C) Top panel: Precision of alignment of imaging and uncaging laser scanning. The horizontal line was bleached with the imaging laser and the spots were bleached with the uncaging laser. Note that the first uncaging spot is precisely found on the „imaging line”. Bottom panel: Precision of uncaging laser spot positioning. 10 uncaging spots were placed at 1 µm intervals. For analysis gaussians were fitted to the bleaching profiles with sub-pixel resolution to obtain their position accurately. Distance between spots is displayed in B).

**Suppl. Figure 4:**
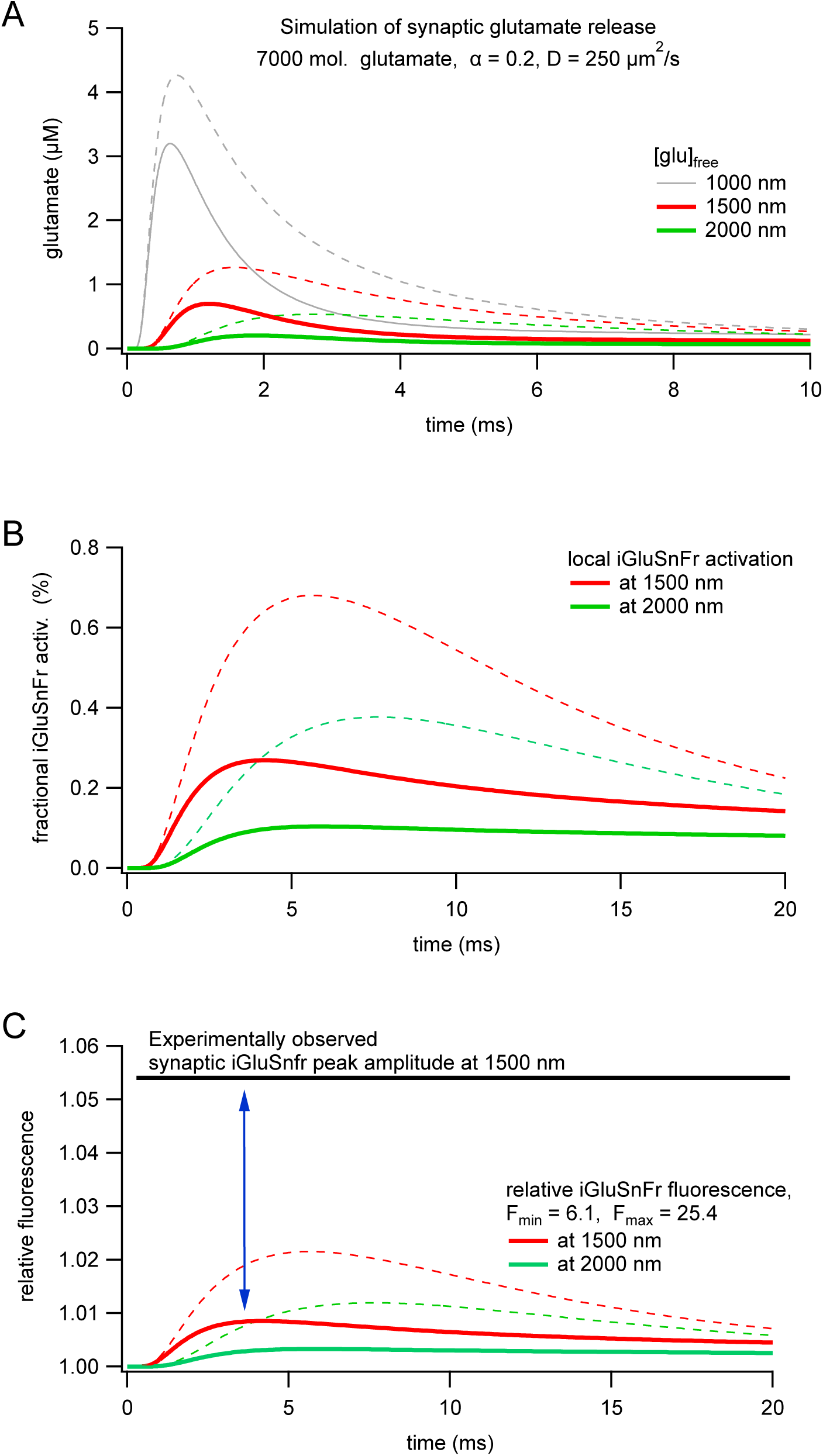
Modeling iGluSnFr activation following synaptic glutamate release - comparison to experiment. (A)The simulation environment c“calc” (*V. Matveev, A. Sherman, R. S. Zucker, Biophysj. 83, 1368– 1373 (2002)*) (as “spherical symmetry”) was used to replicate the neuropil diffusion models of (*D. A. Rusakov, D. M. Kullmann, J. Neurosci. 18, 3158–3170 (1998), B. Barbour, Journal of Neuroscience. 21, 7969–7984 (2001)*) and to simulate synaptic glutamate release, binding to iGluSnFr and fluorescence activation of iGluSnFr. A gaussian-shaped, fusion pore-like source of glutamate (FWHM 5 nm) was placed at the origin and glutamate was released in a peak-like fashion according to: t * sigma^2 * exp(-sigma*t) (t in ms, sigma=39), following (*D. A. Rusakov, D. M. Kullmann, J. Neurosci. 18, 3158–3170 (1998)*). Glutamate transporters were modeled by omitting the translocation step as fixed glutamate “buffers” at a concentration of 100 µM in the extracellular volume. Omitting the translocation is justified as it is slow and has a negligible effect on free glutamate (not shown, also see B. Barbour, Journal of Neuroscience. 21, 7969– 7984 (2001)). No pre- or postsynaptic structures around the release were modeled as previous studies showed that at the distances considered here (>1000 nm) they have almost no effect on the glutamate concentration. Continuous lines throughout this figure represent calculations in the presence of glutamate “buffers” (k+ = 5e06 /(Ms), k- = 100/s), dashed lines in their absence. An effective glutamate diffusion coefficient D=250µm^2/s was employed to account for the tortuousity of the neuropil similar to (*D. A. Rusakov, D. M. Kullmann, J. Neurosci. 18, 3158–3170 (1998), B. Barbour, Journal of Neuroscience. 21, 7969–7984 (2001)*). The vesicle contained 7000 molecules of glutamate according to recent estimates (see discussion for details). Note that at distances of ≥1500 nm free glutamate concentrations remain below 1 µM, show a slowed rise and peak with at least 1 ms delay. (B) The fraction of iGluSnFr molecules reaching the fluorescent state after synaptic release at 1500 and 2000 nm distance from the release site remained below 0.3%. In other words, classical neuropil diffusion models predict minimal iGluSnFr responses. We modeled iGluSnFr according to (*M. Armbruster, C. G. Dulla, J. S. Diamond, eLife. 9, 10404–26 (2020)*) with 3 states: no glutamate bound, glutamate bound and non-fluorescent and glutamate bound and fluorescent. Rate constants were also taken from that study. (C) Conversion of fractional sniffer activation to fluorescent signals using the fluorescence constants for activated and non-activated iGluSnFr molecules indicated (taken from (*M. Armbruster, C. G. Dulla, J. S. Diamond, eLife. 9, 10404–26 (2020)*)). Note that the predicted DF/F iGluSnFr signals remain below 1%, whereas we experimentally determined iGluSnFr amplitudes at a distance of 1500 nm to be ∼5.4% (black horizontal line) following spontaneous, putative quantal, release events. The blue arrow denotes the almost 5-fold differences between the experimental observation and theoretical prediction.

**Suppl. Figure 5:**
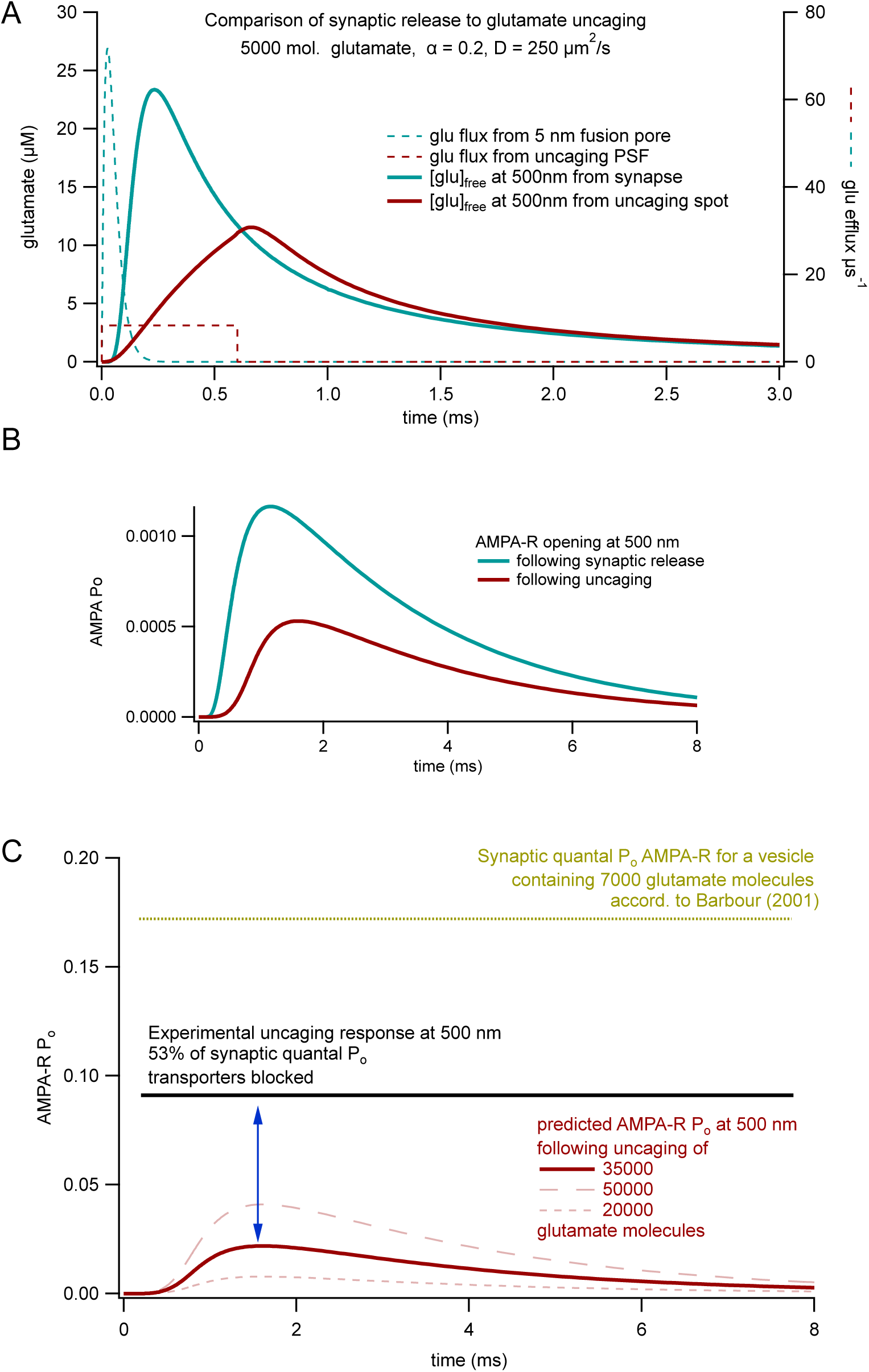
Modeling AMPA-R activation following glutamate uncaging - comparison to experiment. (A)As above the simulation environment “calc” (*V. Matveev, A. Sherman, R. S. Zucker, Biophysj. 83, 1368–1373 (2002)*) was used to replicate neuropil diffusion here in “cylindrical” mode. Neuropil and diffusion properties as in Suppl Fig. 4. This panel compares synaptic glutamate release from a gaussian-shaped, fusion pore-like source of glutamate (FWHM 5 nm) to glutamate released from a PSF. The dashed lines indicate the flux/release rate of glutamate. In this panel, we simulated the release of 5000 glutamate molecules to facilitate comparison to previous work (*B. Barbour, Journal of Neuroscience. 21, 7969–7984 (2001))*. The peak-like release is identical to the one used for Suppl. Fig. 4, 0.6 ms of constant glutamate release was assumed for glutamate uncaging. The source of uncaging glutamate release was assumed to be a 3D gaussian with an x-y FWHM of 280 nm and a 3.5-fold larger FWHM in the z dimension. The blue and red traces illustrate the resulting free glutamate concentration at 500 nm distance. Note that after synaptic release free glutamate rises much quicker and achieves a substantially higher peak glutamate concentration when compared to continuous 0.6 ms release during uncaging. (B) AMPA-R opening is much lower following uncaging release of 5000 molecules of glutamate. AMPRA-R opening was modeled at a distance of 500 nm according to (*D. A. Rusakov, D. M. Kullmann, J. Neurosci. 18, 3158–3170 (1998), B. Barbour, Journal of Neuroscience. 21, 7969– 7984 (2001))*. AMPA-R open probability following synaptic release compares well to (*B. Barbour, Journal of Neuroscience. 21, 7969–7984 (2001)* Fig. 4 therein). (C) Experimentally observed AMPA-R open probabilities in response to uncaging exceed modeling prediction by at least a factor of four. The yellow dashed line indicates the open probability of synaptic AMPA-Rs according to (*B. Barbour, Journal of Neuroscience. 21, 7969–7984 (2001)*). Note that this represents a lower limit when comparing to other modeling studies eg (*D. A. Rusakov, D. M. Kullmann, J. Neurosci. 18, 3158–3170 (1998)*). No glutamate transporters were included here. The black line represents the level of AMPA-R activation we experimentally observed following glutamate uncaging at a distance of 500 nm in the presence of tfb-TBOA. The red line shows in comparison the modeled AMPA-R opening following PSF-shaped uncaging release of 35000 molecules of glutamate (calibrated uncaging content, ∼5 vesicles). Even the release of 50000 molecules of glutamate would be predicted to cause a substantially lower AMPA-R activation (red dashed line). The blue arrow denotes the 4-fold differences between the experimental observation and theoretical prediction.

